# A small protein essential for flagellar assembly and virulence of *Campylobacter jejuni*

**DOI:** 10.1101/2025.04.02.646747

**Authors:** Mona Alzheimer, Kathrin Froschauer, Sarah L. Svensson, Evelyne Hopp, Tina Drobnič, Louie D. Henderson, Deborah A. Ribardo, David R. Hendrixson, Thorsten Bischler, Morgan Beeby, Cynthia M. Sharma

## Abstract

*Campylobacter jejuni* is the leading cause of bacterial food-poisoning, with motility being an essential virulence factor. While many aspects of flagella biogenesis have been studied, a complete picture of the components and regulators of these multi-protein machines is still missing. To identify genes crucial for *C. jejuni* pathogenesis and motility, we applied transposon sequencing in a humanized tissue model. This revealed three largely uncharacterized genes (*pflC*, *pflD*, *pflE*) as essential for motility. While PflC/D turned out to be components of the flagellar motor disk structures, PflE is a small protein of only 57 aa. PflE strikingly affects motor biogenesis, with complete loss of motor structures upon its deletion. We demonstrate PflE is a lipoprotein and supports outer-membrane localization of the main basal-disk protein FlgP. With motility as a critical *Campylobacter* virulence factor, our work demonstrates that deletion of a small protein can bring a bacterial pathogen to a halt.

## Introduction

Despite substantial progress made in understanding how bacterial pathogens establish infection, many molecular factors underlying pathogenesis remain unknown. It is increasingly recognized that core physiological processes, such as gene regulation, metabolism, and especially motility, are integral to virulence of bacterial pathogens (Holden et al., 2004; Rogers et al., 2023). This is particularly true for the zoonotic pathogen *Campylobacter jejuni*, which is the most common cause of bacterial gastroenteritis worldwide (Burnham and Hendrixson, 2018; Havelaar et al., 2015). Despite its prevalence, *C. jejuni* lacks many of the classical virulence factors employed by other enteric pathogens, such as dedicated toxins and secretion systems (Gundogdu et al., 2007; Parkhill et al., 2000). Instead, it relies extensively on motility, surface carbohydrates, cell shape, and metabolic adaptability to establish and cause disease (Burnham and Hendrixson, 2018; Frirdich et al., 2014, 2012; Gao et al., 2017, 2014; Guerry, 2007; Hofreuter, 2014; Liu et al., 2012).

Among these, motility is one of the most central determinants for *C. jejuni* to establish infection. The bacterium’s bipolar flagella enable rapid swimming through viscous environments, and are driven by one of the most complex bacterial flagellar motors characterized to date (Drobnič et al., 2025). The *C. jejuni* motor can generate three times higher torque than that of, e.g., *Salmonella*. This is mediated by elaborate periplasmic disk structures (basal, medial, proximal), which support incorporation of more stator complexes, leading to a wider rotor (C-ring) that increases torque (Beeby et al., 2016; Chaban et al., 2018). While several unique protein components of these structures have been identified (Beeby et al., 2016; Cohen et al., 2024; Feng et al., 2026; Gao et al., 2014), there are likely still unidentified factors involved in flagellar biogenesis, particularly those overlooked that are specific to Epsilonproteobacteria.

One increasingly recognized class of such so far overlooked factors is the one of small proteins (<50 - 100 aa, reviewed in (Gray et al., 2022)), which are notoriously under-annotated and poorly characterized. Once dismissed as translational noise or annotation artifacts, small proteins are now emerging as critical regulators and effectors of bacterial physiology including virulence and stress responses (Burton et al., 2024). Small proteins remain challenging to study due to their small size, biochemical properties, and often absent phenotypes for mutant strains (Yadavalli and Yuan, 2022). However, the development of genomics approaches, peptidomics and especially ribosome profiling approaches has greatly facilitated the discovery of small proteins across phyla (Fuchs and Engelmann, 2023; Gray et al., 2022; Ingolia et al., 2019; Vazquez-Laslop et al., 2022). For example, we recently defined the small proteome of *C. jejuni* (Froschauer et al., 2025) providing a resource to explore the contribution of these elusive factors to key biological processes in bacteria.

To systematically identify novel virulence determinants, including small proteins, we performed a genome-wide transposon (Tn) mutagenesis screen in a tissue-engineered three-dimensional (3D) infection model of the human small intestine (Alzheimer et al., 2020). This model can recapitulate infection phenotypes *in vitro* that had previously only been observed in animal models *in vivo*. Thus, we reasoned that this model might allow the discovery of previously missed factors that contribute to *C. jejuni* fitness in the human intestinal tract, including factors that impact motility.

Our screen uncovered, among others, three genes crucial for infection that turned out to be essential for motility and flagellar motor assembly. We demonstrate Cj0892c (PflD), Cj1643 (PflC), and Cj0978c (PflE) are indispensable for flagellar rotation but not filament biogenesis. Cryo-electron tomography (cryoET) reveals that PflC and PflD are components of the flagellar disks required for stator incorporation. Strikingly, deletion of the gene encoding the small lipoprotein PflE (57 aa, processed length 41 aa) resulted in loss of all disk structures. Our results suggest that, while it is not a structural component of the motor, PflE is required for localization of the major basal disk protein FlgP to the outer membrane, which is the first, essential, step of disk assembly. Our findings broaden our understanding of the complex *C. jejuni* flagellar machinery, an essential virulence determinant and model for studying flagellar function and evolution. Moreover, we highlight PflE as a striking example of a small protein with an essential role in assembling one of the most complex macromolecular machineries in bacteria.

## Results

### Tn-seq in a 3D intestinal tissue model reveals genes required for *C. jejuni* virulence and motility

To screen for *C. jejuni* factors required for host cell interaction in a humanized three-dimensional (3D) tissue model ((Alzheimer et al., 2020), we established a Tn5 transposon mutant pool in the commonly used lab strain NCTC11168 (**Figure S1A**). Identification and quantification of insertion sites by deep sequencing revealed ∼35,000 unique insertions distributed across the ∼1.6 Mbp *C. jejuni* genome (average 1/47 base pairs) (**Figure S1B, Table S1**). The validity of the mutant pool was supported by the lack of insertions in known essential genes (e.g., *cprR*/*Cj1227c*; (Raphael et al., 2005; Svensson et al., 2009)), despite many insertions in adjacent non-essential genes such as *htrA*/*Cj1228c* (Brøndsted et al., 2005) (**Figure S1C, Table S1**). No insertions were identified in 554 protein-coding genes, which largely overlapped with a compilation of putative *C. jejuni* essential genes identified in other studies (S. P. de Vries et al., 2017; Gao et al., 2014; Mandal et al., 2017; Metris et al., 2011; Stahl and Stintzi, 2011).

To identify bacterial genes affecting host interactions, we infected our intestinal 3D tissue model and in comparison classical 2D Caco-2 monolayers with the *C. jejuni* Tn pool (**Figure 1A**). We isolated colony forming units (CFUs) after 4 and 24 hrs of infection in 2D and 3D, respectively, which ensures comparable infection rates (Alzheimer et al., 2020). This included recovery of adherent and internalized bacteria together (ADH), internalized bacteria only after gentamicin treatment (INT), and non-cell-associated bacteria from infection supernatants (SUP). After deep sequencing of Tn5-chromosome junctions (**Table S2**), we identified genes with insertion read count ratios significantly different between SUP and ADH/INT bacteria (**Table S3**; ≥ 10 reads in SUP libraries, log_2_FC ≥ |1|, p ≤ 0.05). This revealed 69 (3D) and 64 (2D) genes where Tn insertion counts significantly decreased for ADH and/or INT vs. SUP, with surprisingly little overlap of 10 genes between the infection models (**Figure 1B**). Genes in this overlap, as well as those uniquely found in 3D, were significantly enriched for the flagellar assembly pathway while no significant functional enrichment was found for genes present in 2D only (**Figure S2**). This underscores the vital role of motility for *C. jejuni* pathogenesis observed in Tn-seq screens in various strains and infection models (S. P. W. de Vries et al., 2017; S. P. de Vries et al., 2017; Gao et al., 2017, 2014; Javed et al., 2010; Johnson et al., 2014; Novik et al., 2010).

**Figure 1.**
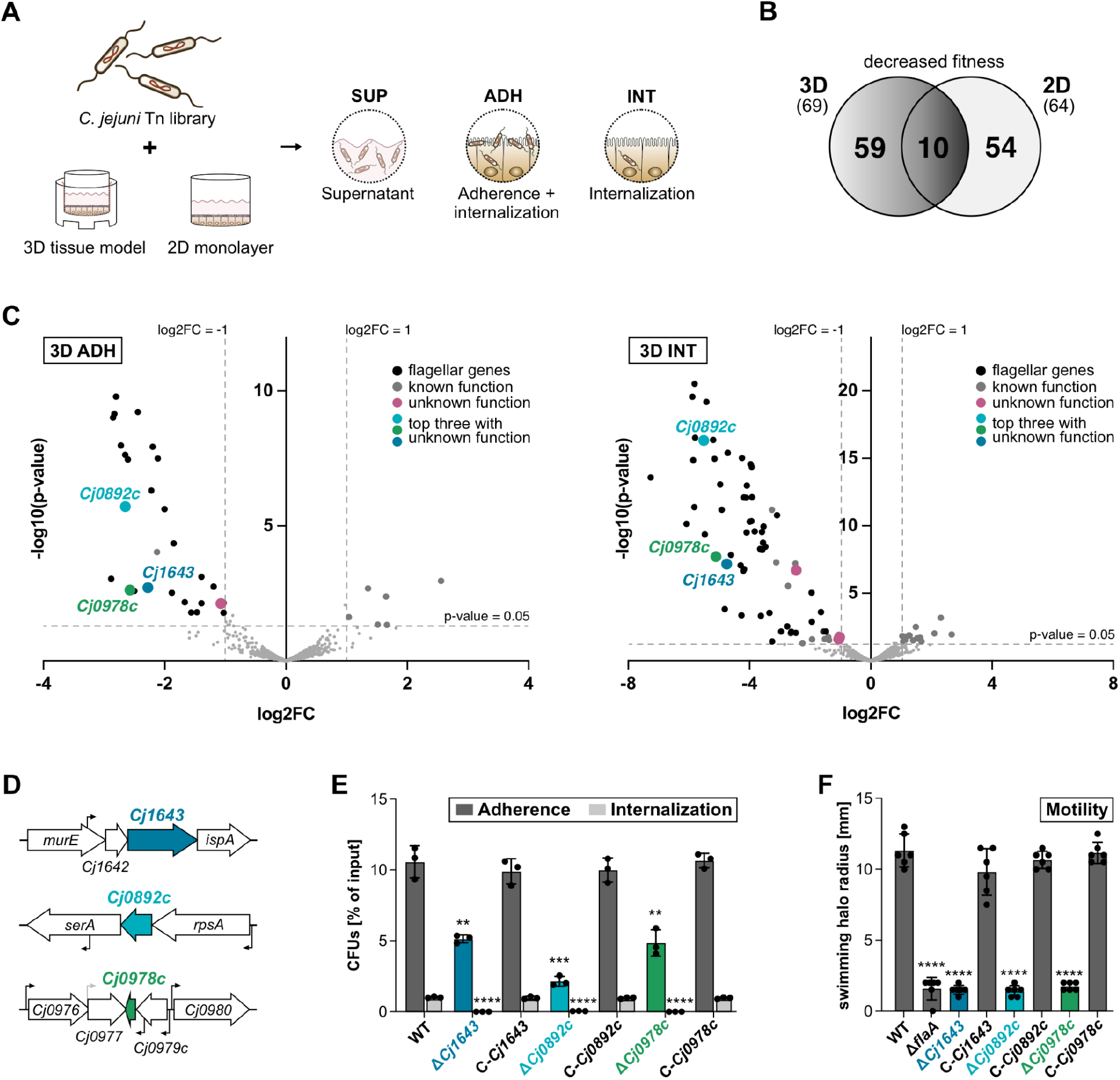
A *C. jejuni* Tn-seq screen in a Caco-2 3D infection model uncovers new factors required for virulence and motility. **(A)** 3D or 2D Caco-2 infection models were infected with a *C. jejuni* transposon (Tn) mutant library. After 24 hrs in 3D and 4 hrs in 2D, CFUs were isolated. SUP: non cell-associated bacteria from the supernatant. ADH: adherent + internalized bacteria. INT: internalized bacteria after an additional 2 hrs of gentamicin treatment. **(B)** Venn diagram depicting the number of genes whose disruption by Tn5 insertion resulted in decreased fitness (ADH+INT) in either the 3D tissue model, the 2D Caco-2 monolayer, or those shared between the two infection systems. Decreased fitness is defined as a significant (p ≤ 0.05) log2FC ≤ ‡1 of gene-wise transposon insertion counts obtained from libraries prepared from adherent and internalized samples (ADH) or internalized samples only (INT) versus those from the supernatant bacteria (SUP; ≥ 10 reads). **(C)** Volcano plots depicting log2FC of insertion counts (x-axis) and -log10 of p-value (y-axis) for *C. jejuni* genes identified in ADH (left) or INT (right) samples versus SUP in 3D tissue model infections of two independent replicates. Horizontal lines mark a p-value of 0.05 and vertical lines a log2FC of |1|. Black = flagellar genes, grey = genes with known function other than motility, pink = genes with thus far unknown function, blue/green = top 3 genes with unknown function. **(D)** Genomic context and transcriptional start sites/operon structure for *Cj1643*, *Cj0892c*, and *Cj0978c* in *C. jejuni* NCTC11168. Bent arrows: TSS (Dugar et al., 2013). **(E)** Validation of Tn-seq infection phenotypes for *Cj1643*, *Cj0892c*, and *Cj0978c* mutants in 3D infection models. CFUs were isolated for adherent and internalized bacteria (Adherence) or internalized bacteria only (Internalization) of NCTC11168 wildtype (WT), deletion (Δ), and complementation (C) strains after 24 hrs p. i. CFUs are displayed as a percentage of input bacteria and represent the mean of three biological replicates with corresponding standard deviations (SDs). Significance is given with respect to the parental WT. **(F)** Soft agar swimming assays for *C. jejuni* NCTC11168 WT and *Cj1643*, *Cj0892*c, and *Cj0978c* deletion (Δ) / complementation (C) strains. Δ*flaA*: non-motile control. Mean of three independent replicates with SD. Significance vs. WT. ****: *p* < 0.0001, ***: *p* < 0.001, **: *p* < 0.01 (Student’s *t*-test).

In line with the important role of motility, the most strongly decreased genes in the 3D tissue model encoded flagellar components (**Figures 1C**). The top three thus-far uncharacterized genes with the largest log_2_FC for decreased fitness for either ADH or INT libraries encoded two putative periplasmic proteins (Cj1643, Cj0892c) and the putative small protein Cj0978c (57 aa) (**Figures 1C**; **Table S3**). While *Cj1643* and *Cj0892c* are likely co-transcribed with their upstream gene(s), *Cj0978c*, which we previously validated as a translated “small protein” (Froschauer et al., 2025), is transcribed from its own promoter located in the upstream gene *Cj0979c* (**Figure 1D**; (Dugar et al., 2013)). Clean deletion mutants in all three genes validated their ADH and INT phenotypes in both infection models (**Figure 1E**; **Figure S3A**). These phenotypes were fully rescued by complementation *in trans* from the unrelated *rdxA* locus. Moreover, we noticed that the deletion mutants in all three genes were completely non-motile in strain NCTC11168 (**Figure 1F**), used for the screen, as well as the widely-used strain 81-176 (**Figure S3B**). Also motility could be restored to WT levels upon complementation in trans (**Figure 1F**). Since flagella and motility are important cell adherence and internalization factors for *C. jejuni*, loss of these functions is likely responsible for the strong infection defects of these mutants.

### *Cj1643*, *Cj0892c*, and *Cj0978c* deletion paralyzes flagella and prevents motor disk assembly

Despite their motility phenotypes, none of the three candidates is encoded near motility-related genes (**Figure 1D**) or regulated by the flagellar sigma factors RpoN or FliA (**Figure S4A**). However, like other flagella-related proteins, such as FliF, FlgP/Q, or PflA/B, all three are conserved in all motile Campylobacterota species inspected (**Figure S4B**; **Table S4**). This suggested that Cj1643, Cj0892c, and Cj0978c play a dedicated role in *C. jejuni* motility and we sought to further decipher the underlying mechanism.

Bacterial motility phenotypes are typically caused by absence of filament expression or defects in motor function. Thus, we first examined the presence of flagellar filaments. Transmission electron microscopy (TEM) of *Cj1643*, *Cj0892c*, and *Cj0978c* deletion and complementation mutants showed that the mutants have WT-like flagella at both cell poles despite their non-motile phenotype (**Figure 2A**). In contrast, a Δ*flaA* control strain expresses only a stubby filament composed of the minor flagellin FlaB (Dugar et al., 2016) (**Figure 2A**). This indicated that Δ*Cj1643*, Δ*Cj0892c*, and Δ*Cj0978c* (hereafter *pflC*, *pflD*, and *pflE*, respectively, for <u>p</u>aralyzed <u>fl</u>agellum) have paralyzed flagella and that their encoded proteins affect motor assembly and/or function. For PflC and PflD, this was supported by a predominantly polar subcellular expression and potential flagella co-localization, determined using strains with C-terminal sfGFP fusions, which are motile (**Figures 2B**; **Figure S4C**). PflE, however, was visible across the whole cell body suggesting an indirect involvement of the protein in flagellar motor function/biogenesis (**Figure 2B**).

**Figure 2.**
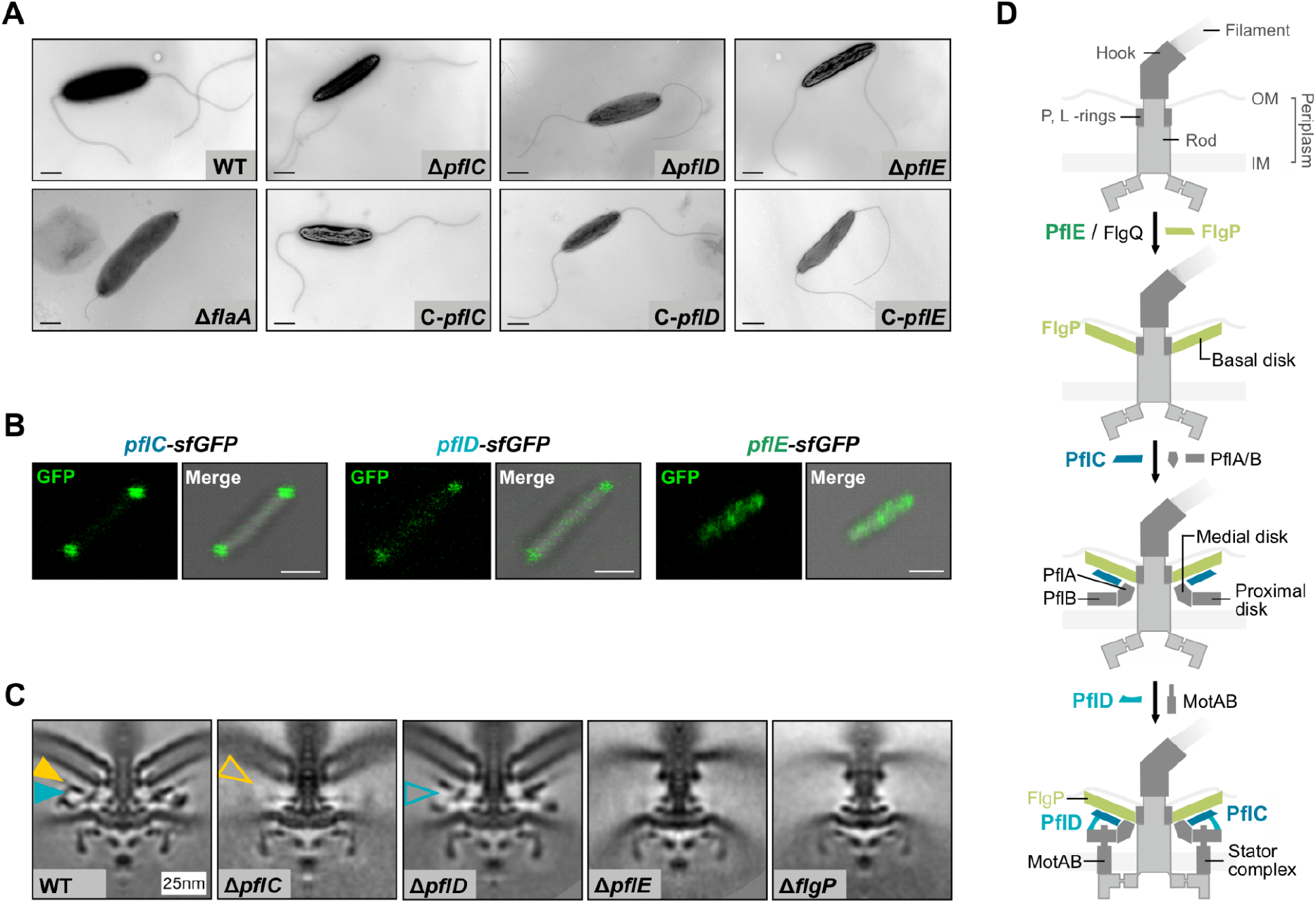
PflC, PflD, and PflE are required for assembly of flagellar motor disk structures. **(A)** Transmission electron microscope (TEM) micrographs of wildtype (WT), Δ*flaA* (lacking most of the flagellar filament), and deletions (Δ) as well as complementations (C) of motility candidates (*Cj1643* = *pflC*, *Cj0892c* = *pflD*, and *Cj0978c* = *pflE*). Bacteria were stained with 2 % uranyl acetate and inspected using a Zeiss EM10 TEM. Images are representative results of two independent experiments. Scale bar: 0.5 µm. **(B)** Confocal microscopy images of *C. jejuni* expressing C-terminally sfGFP-tagged motility candidate proteins. Images are representative of three biological replicates. Scale bar: 1 µm. **(C)** Left to right: 100 nm x 100 nm slices through subtomogram averages of wild-type (from EMD-3150), Δ*pflC*, Δ*pflD*, Δ*pflE*, and Δ*flgP* (from EMD-3159) flagellar motors. PflC is likely to be the large lattice that disappears upon deletion of *pflC* as indicated by yellow arrowhead for presence (filled) and absence (empty); PflD is likely to be a peripheral post-like structure that disappears upon deletion of *pflD* as indicated by cyan arrowhead for presence (filled) and absence (empty). Location of PflE is less unambiguous, but the basal disk formed of FlgP cannot assemble upon its deletion. **(D)** Overview of flagellar disk assembly in *C. jejuni* (adapted from (Beeby et al., 2016)) and cartoon representation of the *C. jejuni* motor with location/importance of PflC, PflD, and PflE based on this work and (Beeby et al., 2016; Drobnič et al., 2025). OM: outer membrane, IM: inner membrane.

To investigate the involvement of PflC, -D & -E in the flagellar motor further, we imaged the motor structures of *C. jejuni* NCTC11168 Δ*pflC*, Δ*pflD*, and Δ*pflE* mutants by cryo-electron tomography (cryoET) and subtomogram averaging. Comparison to a WT strain suggested that PflC is a novel component of the medial disk, which assembles immediately below the basal disk, and that PflD forms a density on the periphery of the proximal disk and is required for incorporation of stator complexes (**Figure 2C**). This, we recently further confirmed in a high-resolution in situ structure of the *C. jejuni* motor (Drobnič et al., 2025). Surprisingly, deletion of *pflE* rendered the motor completely devoid of all periplasmic structures, while the core axial components were still assembled (**Figure 2C**). This phenotype mimicked motor structures imaged for Δ*flgP* (**Figure 2C**) or Δ*flgQ* (Beeby et al., 2016) deletion mutant strains. Primarily, the Δ*pflE* motor lacked the basal disk composed of FlgP, while other missing periplasmic components were likely absent because they require a basal disk for assembly (Beeby et al., 2016). While we could not assign the small protein as a structural component of the flagellar motor, our findings suggested that PflE is an essential component in the *C. jejuni* basal disk assembly pathway outlined in the following updated model (**Figure 2D**): FlgP first polymerizes as the basal disk with the help of PflE and FlgQ, the latter of which is known to be required for FlgP stability and/or localization (Beeby et al., 2016; Drobnič et al., 2025; Henderson and Beeby, 2018; Henderson et al., 2020; Sommerlad and Hendrixson, 2007). PflC then assembles as a component of the medial disk, providing a platform for PflA and PflB, which together with the stator complexes, forms the proximal disk. Finally, integration of the stator units seems to require PflD. Thus, while PflC and PflD have direct structural roles in the *C. jejuni* flagellar motor (Drobnič et al., 2025), PflE might support motor assembly indirectly.

### The small protein PflE affects FlgP expression and localization

It is remarkable that a small protein of only 57aa has such a large impact on a complex macromolecular machine. Small proteins often act via direct interactions with other proteins or protein complexes, and regulate, localize, or support their function (reviewed in (Gray et al., 2022)). Since the motor structure of Δ*pflE* phenocopied that of Δ*flgP* and Δ*flgQ* (**Figure 3A**; **(Beeby et al., 2016)**), we investigated the effect that these three proteins have on each other. Deletion of neither *flgP* nor *flgQ* affected protein levels of 3xFLAG-tagged PflE (**Figure 3B**, left). Subcellular fractionation combined with western blot showed that PflE mainly localizes to the inner membrane and the cytoplasm with a smaller fraction detectable in the outer membrane (**Figure 3B**, right). Deleting *flgP* or *flgQ* had no effect on PflE localization, independent of the *C. jejuni* isolate tested (**Figure 3B**, right; **Figure S5A** for strain NCTC11168; **Figure S5B** for strain 81-176). Thus, FlgP and FlgQ neither influence PflE protein levels nor its subcellular localization (**Figure 3B**, lower). In contrast, deletion of *pflE* increased FlgQ-3xFLAG protein levels two-fold, while deletion of *flgP* led to a 2.5-fold decrease in FlgQ levels (**Figure 3C**, left; (Beeby et al., 2016)). In a double deletion mutant of *pflE* and *flgP*, FlgQ levels were decreased comparable to that of Δ*flgP* alone, suggesting that the influence of FlgP on FlgQ takes precedence over that of PflE. Subcellular fractionation detected FlgQ mainly in the cytoplasmic and periplasmic fractions, which did not change upon deletion of *pflE* (**Figure 3C**, middle; **Figure S5C**). Thus, while PflE slightly increased FlgQ levels, it seems to have no effect on its subcellular localization (**Figure 3C**, right).

**Figure 3.**
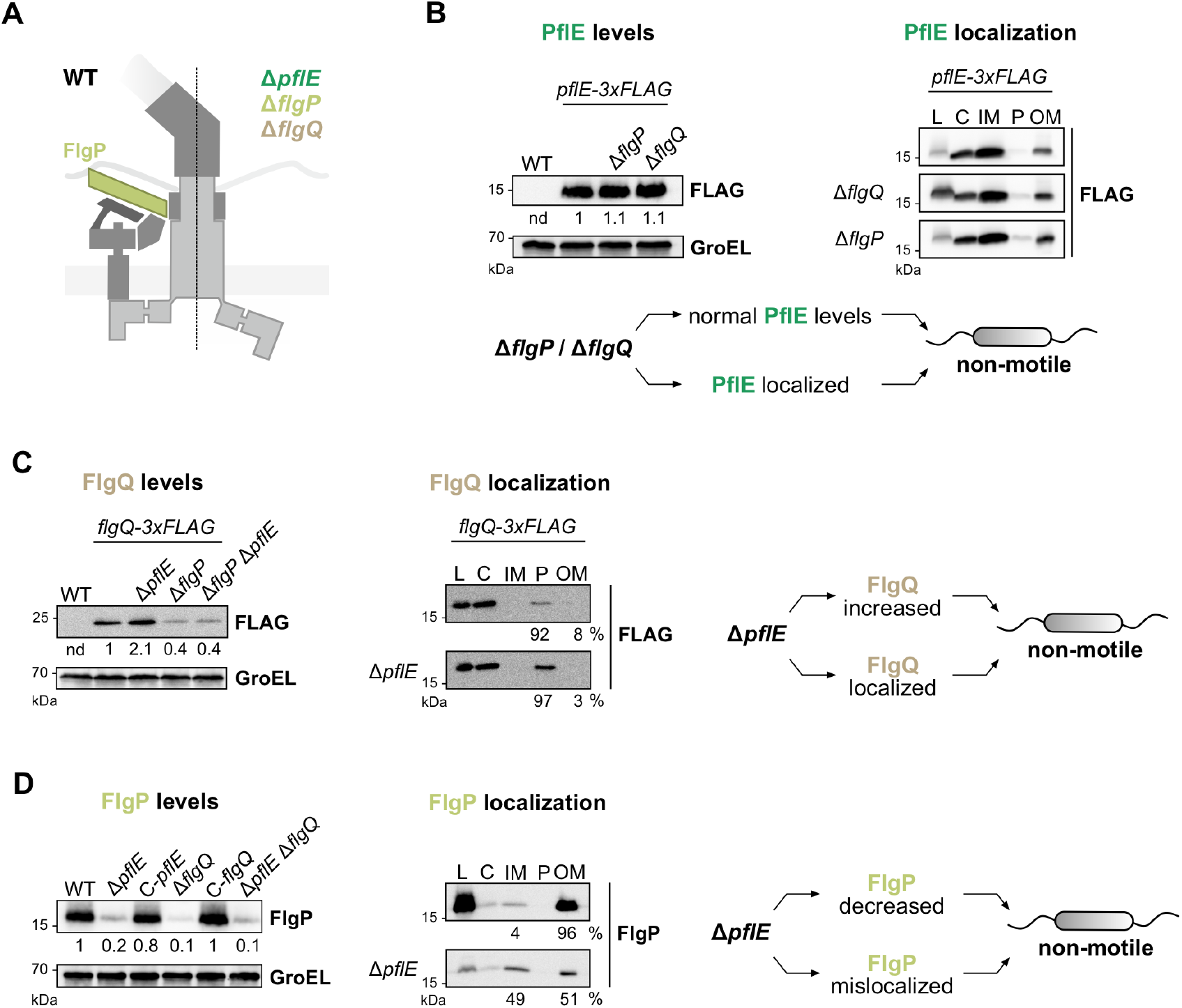
PflE affects FlgP localization and stability. **(A)** Schematic representation of cryoET structures present in the *C. jejuni* wild-type motor (WT, left) and deletion (Δ) of *pflE*, *flgP*, or *flgQ* (right). **(B-D)** PflE-3xFLAG **(B)**, FlgQ-3xFLAG **(C)**, FlgP **(D)** protein levels and subcellular fractionation either in wild-type or Δ*flgP*/*flgQ*/Δ*pflE* backgrounds analyzed by western blot. Summary cartoons for each panel are depicted below **(B)** or on the right **(C, D)**. PflE and FlgQ were detected with anti-FLAG and FlgP with a specific antiserum (Sommerlad and Hendrixson, 2007). GroEL served as loading control. L: lysate; C: cytoplasm; IM: inner membrane; P: periplasm; OM: outer membrane. Bands were quantified with Aida software (Raytest) and are relative values compared to wild-type backgrounds for protein levels (left). nd: not determined. For quantification of subcellular fractions **(C**, **D**, middle), values represent percentages relative to the combined band intensity of periplasmic and outer membrane fraction **(C)** or inner and outer membrane fraction **(D)**. For subcellular fractionation controls see Figure S5.

Lastly, we investigated the effect of PflE on the basal disk component FlgP and found that FlgP levels were almost completely abolished by deletion of the small protein, similar as in a Δ*flgQ* mutant and could be restored to WT levels upon complementation (**Figure 3D**, left; (Beeby et al., 2016)). Since levels of FlgP in a double deletion of *pflE* and *flgQ* did not decrease further, this suggests that they both affect FlgP in the same pathway. To determine how PflE might affect FlgP levels, we tested *flgP* transcriptional and translational reporter fusions, and found that *pflE* had no effect on *flgP* transcription or translation (**Figure S6A**, **S6B**). While PflE had no major or only a minor effect on *flgP* transcript stability (**Figure S6C**), we found that FlgP protein was less stable in Δ*pflE* (**Figure S6D**). FlgP is predominantly located in the outer membrane (**Figure 3D**, middle; (Sommerlad and Hendrixson, 2007)). We therefore tested the effect of *pflE* on FlgP localization, hypothesizing that mislocalization might reduce protein stability. In line with this, deleting *pflE* shifted the distribution of FlgP from predominantly outer membrane to approximately equally distributed between inner and outer membranes (**Figure 3D**, middle; **Figure S5D**). This reflects the FlgP pattern observed in Δ*flgQ* (**Figure S5A**; (Beeby et al., 2016; Sommerlad and Hendrixson, 2007)). Thus, the small protein PflE dramatically reduces protein levels of FlgP and alters its subcellular localization (**Figure 3D**, right). Together with the observation that neither FlgP nor FlgQ in turn affect localization of PflE, this suggests that PflE acts upstream of FlgP (and possibly FlgQ).

### PflE is a lipoprotein localized to the inner and outer membrane

PflE is annotated as a lipoprotein (Parkhill et al., 2000). Lipoproteins are generally identified based on the presence of a signal sequence and a lipobox motif with one invariant Cys ([LVI][ASTVI][GAS][**C**]), to which the lipid anchor is added (**Figure S7A**). SignalP 6.0 (Teufel et al., 2022) also predicted PflE to be a lipoprotein in *C. jejuni* with the invariant Cys of the putative lipobox (ISG<u>C</u>) at position 17 (**Figure S7B**). Moreover, sequence alignment and signal peptide/lipobox predictions of PflE homologs in Campylobacterota revealed that in particular the invariant Cys (C17 in *C. jejuni* NCTC11168) is fully conserved (**Figure 4A**; **Figure S7C**; **Table S5**). To verify the lipoprotein status of PflE in *C. jejuni*, we substituted the invariant Cys for Ala (M1, **Figure 4A**) and analysed motility, PflE protein levels, and its subcellular localization pattern. Mutation of C17 in PflE caused a complete loss in motility similar to a deletion mutant of *pflE* (M1, **Figure 4B**). Mutation of the invariant Cys also reduced PflE-3xFLAG protein levels by five fold and changed its migration pattern on western blots compared to wild-type PflE, consistent with interference with lipoprotein processing (**Figure 4C**, left). In addition to reduced protein levels, cell fractionation showed that PflE levels in the M1 mutant were drastically reduced in the inner membrane and completely absent from the outer membrane, but instead increased in the periplasm (**Figure 4C**, middle; **Figure S7D**). Taken together, decreased protein levels, subcellular mislocalization, and complete loss of motility of a PflE lipobox mutant (**Figure 4C**, right) strongly indicates that PflE is a *bona fide* lipoprotein present mostly in the inner and to a lesser degree in the outer membrane.

**Figure 4.**
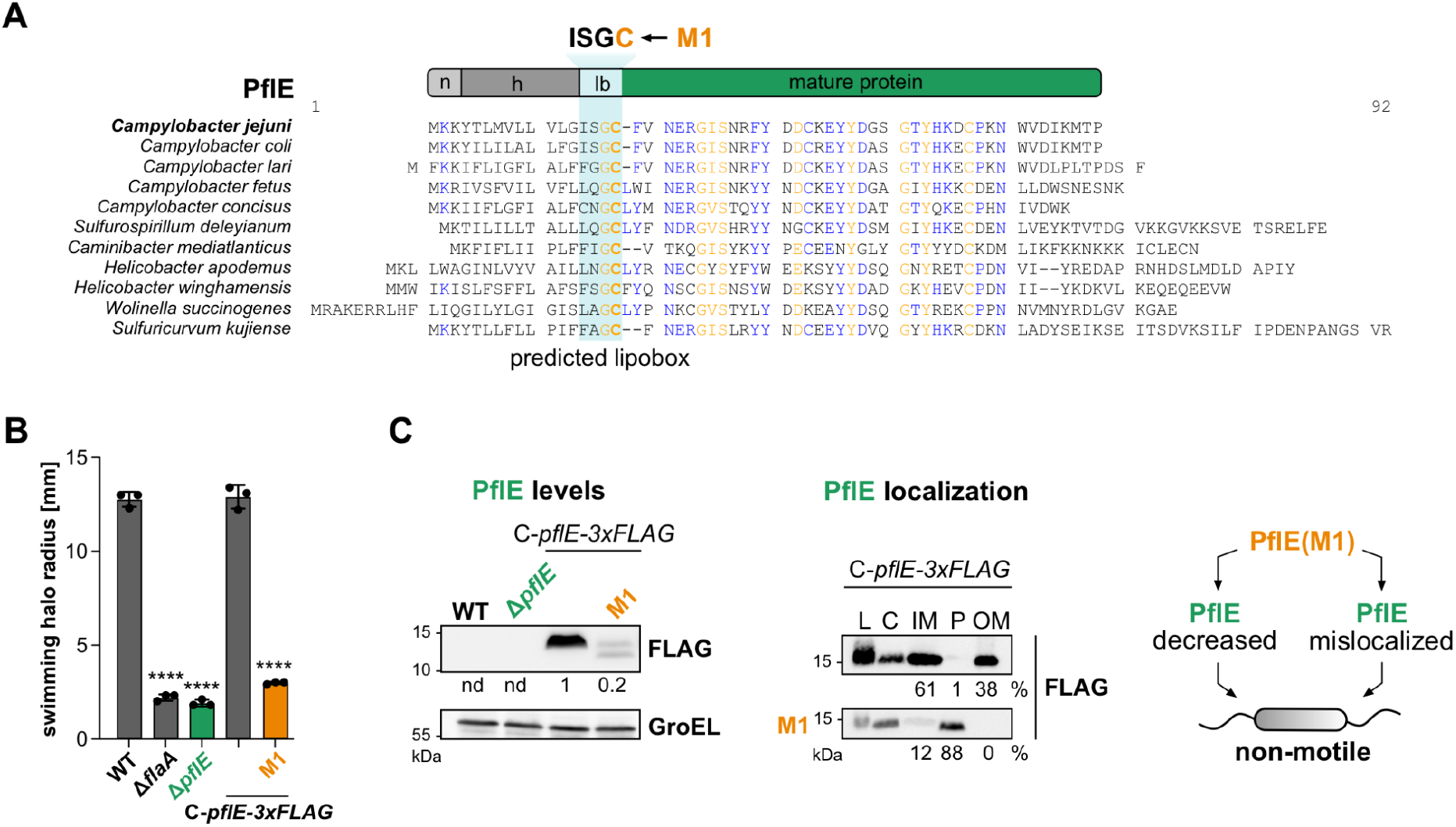
The small protein PflE is a lipoprotein in *C. jejuni*. **(A)** Schematic representation and alignment of PflE protein sequences from indicated strains using Multalin (Corpet, 1988). Putative lipoprotein regions predicted by SignalP – 6.0 (Teufel et al., 2022) (see also Figure S7B, Table S5): n = n-region (positively charged region), h = h-region (hydrophobic region), lb = lipobox. M1 (orange): Conserved invariant cysteine residue of the predicted lipobox sequence exchanged to Ala via site-directed mutagenesis. **(B)** Motility assay of WT, Δ*pflE*, and complementation with PflE-3xFLAG in *trans* (C-*pflE*-3xFLAG) or with Cys-Ala point mutation (M1) in 0.4% soft agar. Δ*flaA*: non-motile control. Error bars: SD of the mean of independent replicates (n = 3). ****: *p* <0.0001 (Student’s *t*-test). **(C)** PflE-3xFLAG protein levels (left) and subcellular fractionation (middle) either in wildtype, Δ*pflE*, or *pflE* M1 point mutant (C-terminally 3xFLAG tagged). Cartoon summarizing the results is depicted on the right. PflE was detected with anti-FLAG. GroEL: loading control. L: lysate; C: cytoplasm; IM: inner membrane; P: periplasm; OM: outer membrane. Bands were quantified with Aida software (Raytest) and are relative values compared to wild-type backgrounds for protein levels. nd: not determined. For quantification of subcellular fractions (middle), values are percentages relative to the combined band intensity of periplasmic, inner, and outer membrane fraction. For fractionation controls see also Figure S7D.

### FlgP lipoprotein status differs in *C. jejuni* and *H. pylori*

The observation that stability and localization of FlgP in *C. jejuni* was affected by the small lipoprotein PflE was surprising, as FlgP itself is annotated as a lipoprotein in *C. jejuni* (Parkhill et al., 2000). SignalP 6.0 (Teufel et al., 2022) predicted both PflE and FlgP to be lipoproteins in *Campylobacter* species, *Sulfuricurvum*, *Nautilia*, and *Caminibacter*, with the invariant Cys of FlgP being generally conserved (**Figure 5A**, **5B**; **Figure S8A**; **Table S5**). Curiously though, a FlgP lipobox was only predicted in *Helicobacter* species lacking a homolog of PflE, while in species encoding the small protein, FlgP lacked a predicted lipobox sequence (**Figure 5A**, **5B**; **Figure S8A**; **Table S4, S5**). Thus, it appears that not all FlgP homologs are lipoproteins and raised the question of whether the *C. jejuni* FlgP lipobox is in fact functional, despite conserving the invariant Cys. To test this, we exchanged the lipobox Cys residue in FlgP of *C. jejuni* (**Figure S8B**, left) and *H. pylori* (**Figure S8B**, right) for Ala (**Figure 5C**, CjFlgP-C17A/HpFlgP-C25A) and measured swimming of both species in soft agar plates. Surprisingly, *C. jejuni* motility assays revealed that the Cys-Ala lipobox mutant of FlgP (CjFlgP-C17A) rescued the motility defect of Δ*flgP* to a similar extent as a complementation with wild-type FlgP (80-90% of the WT strain) (**Figure 5D**, left). In *H. pylori*, the corresponding lipobox mutant (HpFlgP-C25A) was completely non-motile, like a Δ*flgP* mutant (**Figure 5D**, right). Thus, our results indicate that while HpFlgP requires a functional lipobox to be motile, CjFlgP does not seem to be dependent on the presence of the invariant Cys for motility.

**Figure 5.**
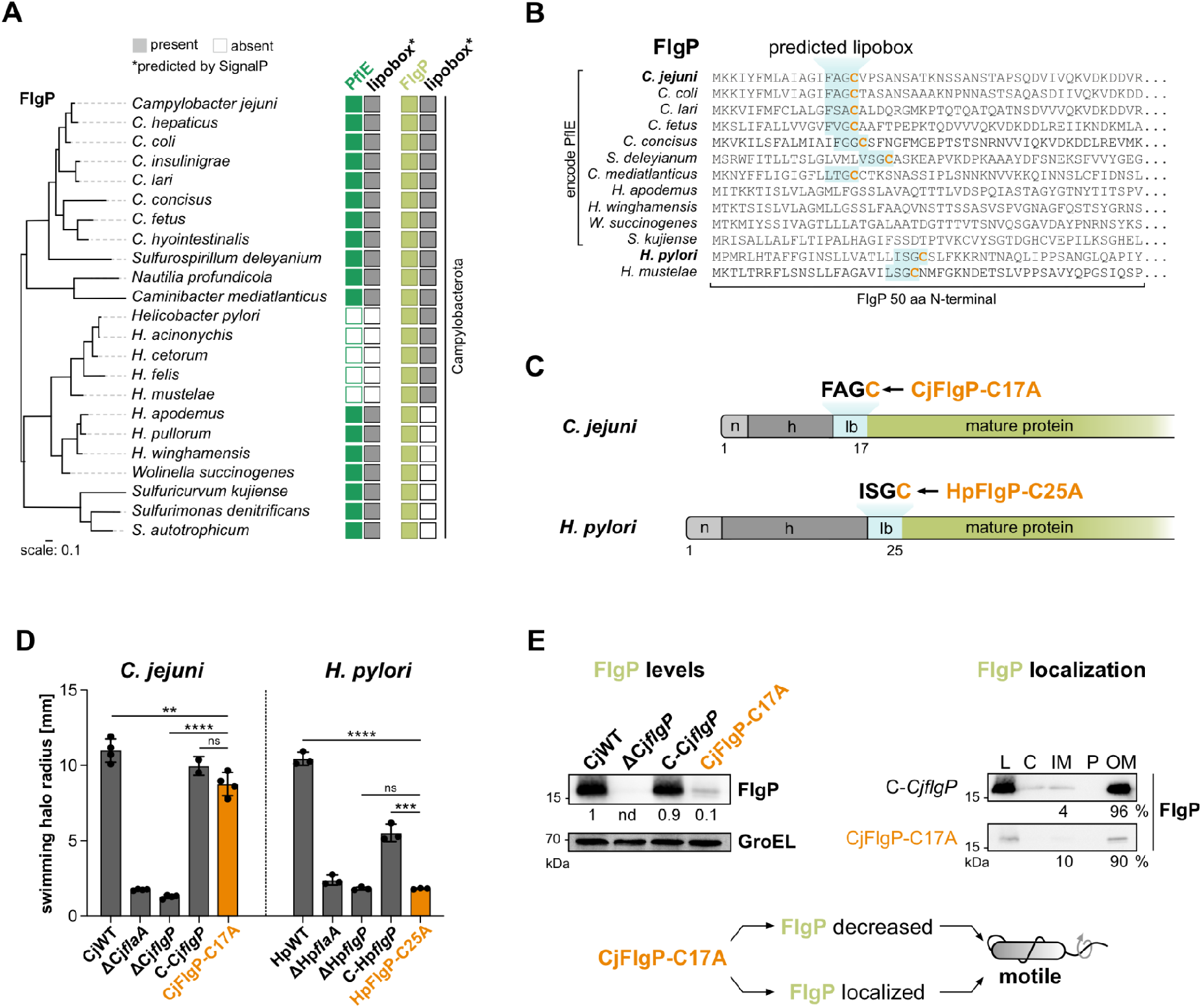
FlgP is not a lipoprotein in *C. jejuni*. **(A)** Conservation analysis of PflE and FlgP in representative Campylobacterota species. Filled boxes: detected (BLASTp)/predicted (Signal P). Species are organized based on a phylogenetic tree based on FlgP (left). See **Methods, Fig. S4B, and Table S4** for additional details. **(B)** 50 N-terminal amino acid (aa) sequences of FlgP orthologs in indicated strains. Lipoboxes were predicted with SignalP – 6.0 (Teufel et al., 2022) (see also Table S5) and are highlighted in blue with the invariant Cys residue marked in bold and orange. **(C)** Schematic representation of native FlgP protein in *C. jejuni* and *H. pylori* with putative lipoprotein regions: n = n-region (positively charged region), h = h-region (hydrophobic region), lb = lipobox. Cys mutations are indicated with an arrow (CjFlgP-C17A for *C. jejuni* and HpFlgP-C25A for *H. pylori*). **(D)** Motility assay demonstrating the effect of FlgP lipobox point mutation for CjFlgP-C17A (left) and pFlgP-C25A (right). Depicted is the mean of three and four independent replicates for *H. pylori* and *C. jejuni*, respectively, with corresponding SDs (for C-Cj*flgP* n=2). ****/***/**: *p* <0.0001/<0.001/<0.01. ns: not significant (Student’s *t*-test). **(E)** *C. jejuni* FlgP protein levels (left) in wildtype (CjWT), Δ*flgP*, and complementation with wild-type FlgP (C-Cj*flgP*) or the lipobox mutant (CjFlgP-C17A), as well as subcellular fractionation (right) of C-Cj*flgP* and CjFlgP-C17A. Lower panel: cartoon summarizing the results of panels D and E. FlgP was detected with FlgP antiserum. GroEL: loading control. L: lysate; C: cytoplasm; IM: inner membrane; P: periplasm; OM: outer membrane. Bands were quantified with Aida software (Raytest) and are relative values compared to wild-type backgrounds for protein levels. nd: not determined. For quantification of subcellular fractions, values are percentages relative to the combined band intensity of inner and outer membrane fraction. For fractionation controls see also Figure S8.

Interestingly, while the *C. jejuni* C17A mutant retained almost WT motility, FlgP protein levels were decreased 10-fold compared to wildtype and the complementation strain (C-*CjflgP*) (**Figure 5E**, left). This is in line with our previous observation that lower FlgP levels can still support *C. jejuni* motility (Cohen et al., 2024). However, subcellular fractionation of CjFlgP-C17A showed that, even though FlgP levels were low, the protein still localized predominantly to the outer membrane as in the WT strain (**Figure 5E**, right; **Figure S8C**). The fact that a *C. jejuni* FlgP lipobox mutant retains motility as well as WT localization despite lower levels of FlgP protein (**Figure 5E**, lower) suggests that in *C. jejuni*, unlike in *H. pylori*, FlgP is not a lipoprotein.

### PflE lipobox function is required for FlgP localization to the outer membrane

Our observation that Δ*pflE* affects FlgP localization, together with the absence of a functional lipobox in FlgP for Campylobacterota that also encode PflE (**Figure 5A**; **Figure S8A**), implicates the small protein, rather than the lipid modification, in FlgP localization. To test this, we measured FlgP protein levels and localization in the M1 lipobox mutant of PflE. Mutating the invariant Cys of PflE phenocopied the reduced FlgP levels of Δ*pflE* (**Figure 6A**, left). Like for the *pflE* deletion mutant, this is likely due to an effect of the PflE-M1 mutation on FlgP protein stability (**Figure S9A**). Likewise, we also observed a shift of FlgP distribution from mainly outer membrane to inner membrane in the PflE lipobox mutant (**Figure 6A**, middle; M1; **Figure S9B**). Thus, mutating the PflE lipobox affected FlgP protein levels and localization (**Figure 6A**, right), indicating that FlgP requires PflE to be a functional lipoprotein for proper localization to the outer membrane.

**Figure 6.**
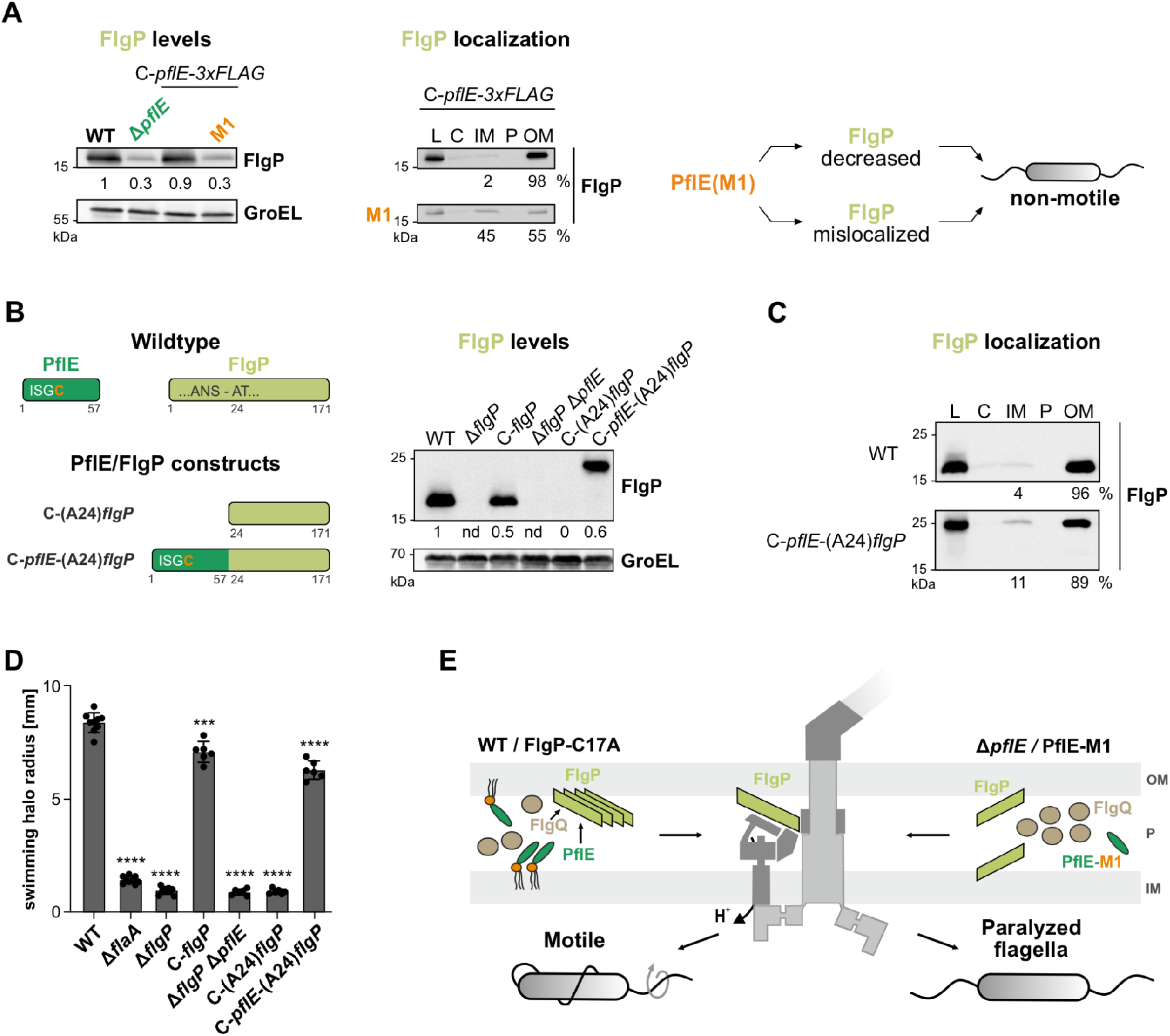
Lipoprotein function of PflE affects FlgP and a PflE-FlgP fusion can restore motility. **(A)** FlgP protein levels (left) and subcellular fractionation (middle) either in wildtype, Δ*pflE*, and complementation with PflE-3xFLAG (C-*pflE*-3xFLAG) or with Cys-Ala point mutation (M1). Cartoon summarizing the results is depicted on the right. FlgP was detected with FlgP antiserum. GroEL: loading control. L: lysate; C: cytoplasm; IM: inner membrane; P: periplasm; OM: outer membrane. Bands were quantified with Aida software (Raytest) and are relative values compared to wild-type backgrounds for protein levels. For quantification of subcellular fractions, values are percentages relative to the combined band intensity of inner and outer membrane fraction. For fractionation controls see also Figure S10B. **(B)** Left: Schematic representation of PflE and FlgP in *C. jejuni* wildtype, as well as FlgP constructs that were expressed from the unrelated *rdxA* locus of a Δ*flgP* Δ*pflE* double deletion mutant. C-(A24)*flgP* is an N-terminally truncated version of FlgP lacking the predicted signal peptide sequence (aa 1-24). C-*pflE*-(A24)*flgP* is a fusion construct of full length PflE (57 aa) fused to the 24^th^ amino acid of FlgP. Right: Western blot analysis of WT, *flgP* deletion (Δ*flgP*) and complementation (C-*flgP*), Δ*flgP* Δ*pflE*, C-(A24)*flgP*, and C-*pflE*(A24)*flgP*. FlgP was detected with FlgP antiserum and GroEL served as loading control. Protein levels were quantified as in (A, left). nd: not determined. **(C)** Subcellular fractionation of *C. jejuni* WT and C-*pflE*(A24)*flgP*. Detection of FlgP with FlgP antiserum. Bands were quantified. Bands were quantified as described for (A, middle). For fractionation controls see also Figure S10B. **(D)** Motility assay of the *C. jejuni* strains in (B). Error bars: SD of the mean of independent replicates (n = 3). ****: *p* <0.0001, ***: *p* <0.001 vs. WT (Student’s *t*-test). **(E)** Working model of the role of PflE in flagellar biogenesis. The small lipoprotein and its membrane association is required for localization of FlgP to the outer membrane and thereby for motility in *C. jejuni*.

To provide additional evidence that FlgP depends on the small protein, we generated a PflE-FlgP fusion protein. Although CjFlgP does not appear to be a lipoprotein, SignalP predicted a potential signal peptidase I (SPI) cleavage site between position S23 and A24 of the FlgP lipobox mutant (**Figure S10A**) indicating the presence of a signal peptide. To exclude any impact of this signal peptide on FlgP localization, we N-terminally fused the 57-aa long PflE to residue A24 of FlgP (C-*pflE*-(A24)*flgP*) (**Figure 6B**, left). As a control, we also generated a strain expressing only the truncated version of FlgP lacking the signal peptide sequence (C-(A24)*flgP*). Both fusion proteins were expressed from the *rdxA* locus of a Δ*flgP* Δ*pflE* double deletion mutant. Analyzing total protein samples via western blot showed that the truncated version of FlgP (C-(A24)*flgP*) was either not expressed or could not be detected by the FlgP antiserum (**Figure 6B**, middle). In contrast, the fusion of PflE and FlgP (C-*pflE*-(A24)*flgP*) was detected at the expected size and showed only slightly lower expression levels compared to wildtype (WT) or a FlgP complementation strain (C-*flgP*). Since it does not appear to be FlgP protein levels but proper FlgP localization to the outer membrane that is crucial for *C. jejuni* motility, we also analyzed subcellular fractionation samples of the FlgP constructs in the Δ*pflE* Δ*flgP* double deletion mutant. While truncated FlgP was not detectable, the PflE-FlgP distribution was similar to that of wild-type FlgP (**Figure 6C**; **Figure S10B**). The truncated FlgP version could not rescue motility of the Δ*flgP* Δ*pflE* double deletion. However, the strain expressing the fusion of PflE-(A24)FlgP was motile, albeit to a lesser extent as the wildtype or a FlgP complementation strain (**Figure 6D**).

Overall, our data point towards a model (**Figure 6E**) where FlgP is a degenerate lipoprotein in *C. jejuni* and does not rely on its own invariant Cys as a lipid anchor for localization in the outer membrane. By comparison, the small protein PflE is a lipoprotein that predominantly localizes to the inner membrane. This membrane association of PflE is crucial for efficient FlgP localization in the outer membrane and, as a consequence, for *C. jejuni* motility.

## Discussion

We have applied Tn-seq during infection of a 3D intestinal tissue-engineered infection model as well as of 2D Caco-2 monolayers to identify *C. jejuni* factors involved in host interactions. This led to the identification of three previously uncharacterized motility factors, PflC, PflD, and PflE, and highlights motility as one of, if not the most predominant virulence determination of this intestinal pathogen. One of the identified genes encodes the small lipoprotein PflE, which appears to be the first small protein with a direct role in flagellar motor biogenesis, making it essential not just for motility but also for pathogenesis of *C. jejuni*.

While small proteins have increasingly been identified across bacterial genomes, their functional characterization remains limited. Our study showed that PflE is a lipoprotein in *C. jejuni* indispensable for outer membrane localization of the flagellar basal disk protein FlgP. Beyond PflE, there are so far virtually no small proteins, other than the anti-sigma factor FlgM (65 aa), with demonstrated roles in motility or flagellar biogenesis, and none that are absolutely required. Recently, three paralogous small proteins in *Salmonella* were linked to motility albeit indirectly via regulation of PhoP (Oguri et al., 2019). Similarly, an *mgrB* deletion mutant in *Salmonella* also showed reduced motility, most likely again an indirect effect via PhoPQ regulation (Venturini et al., 2020). As small proteins are well known to modulate or be part of membrane complexes (Gray et al., 2022), it is surprising that so far no small protein has been directly associated with the flagellar structure. PflE, at only 41 aa long (for its putative processed length), is thus a pioneering example of small proteins that are essential for the structure and function of the flagellum and thereby motility, which is a vital virulence trait for many bacterial pathogens.

The motor structures of the Campylobacterota are more complex compared to those in, *e.g*., *E. coli* or *Salmonella* with many additional accessory structures (Chaban et al., 2018). For two of our newly discovered proteins, PflC and PflD, we recently confirmed in a separate study of a subnanometre-resolution structure of the *C. jejuni* periplasmic motor in situ that they are structural components of the motor, and specifically the medial disk (Drobnič et al., 2025). While PflD and PflE were identified in previous *C. jejuni* screens (but not highlighted or characterized) (S. P. de Vries et al., 2017; Gao et al., 2017), PflC seems to be unique to our dataset. It forms the inner band of the medial disk and connects to the basal disk. PflD, together with the recently characterized flagellar cage proteins FcpMNO, form a peripheral cage structure (Feng et al., 2026). It was proposed that PflD could take on alternating extended and contracted conformations, forming peripheral cage units with FcpMNO in its contracted state while connecting to PflC in the extended form. This cage, in connection with the E-ring (FlgY) and the spoke-rim structure (PflA/B), provides a scaffolding platform that is critical for stator assembly (Feng et al., 2026; “The building blocks of one of the most complex flagellar nanomachines.,” 2026). Interestingly, while the E-ring appears to be of an ancient origin, the cage structure consisting of PflD and FcpMNO seems to be a unique adaptation of the Campylobacterota and shows structural homology to type IV pili components (Feng et al., 2026; “The building blocks of one of the most complex flagellar nanomachines.,” 2026). However, even within Campylobacterota motor structures, several differences between species have evolved. For instance, *H. pylori* and *Wolinella* lack the PflC-corresponding density below the basal disk (Cohen et al., 2024), despite encoding apparent homologs. It will be interesting if they share the same function, as PflC itself might have evolved from an HtrA protease (Drobnič et al., 2025). It remains unclear why these different structures have evolved between the different motor structures, but identifying molecular and functional differences is the first step to understanding this.

We were not able to assign a density to the small lipoprotein PflE in either this study, or the more recent high-resolution structures (Drobnič et al., 2025). While FlgP itself is annotated as a lipoprotein, our findings suggest that, in *C. jejuni*, its outer membrane localization does not depend on the canonical lipobox cysteine, but instead relies on PflE. This indicates that PflE could function as an anchor for CjFlgP at the outer membrane. A comparable anchoring role has, e.g., been described for the small membrane protein Prli42, which anchors the stressosome in *Listeria* to the inner membrane (Impens et al., 2017). If this is the case for PflE, it raises the question of why *C. jejuni* and potentially also other Campylobacterota have retained two separate proteins, compared to, e.g., *H. pylori*, which conserves a functional lipobox in FlgP. In addition, PflE appears to be mainly located in the inner membrane, while the final destination of FlgP is the outer membrane. This suggests that PflE is not an anchor, per se, but responsible for stabilizing FlgP until the assembly of the distal rod as well as L- and P-rings is completed (Beeby et al., 2016). Either of the above-mentioned scenarios would indicate that PflE and FlgP interact with each other. So far we were not able to confidently show interaction between the two proteins, which might be due to their membrane association. *C. jejuni* PflE and FlgP were also tested in a yeast two-hybrid screen (Parrish et al., 2007), but did not show a confident interaction.

Alternatively, PflE could be involved in the shuttling of FlgP from the inner to the outer membrane and therefore interact with it only transiently. If this is the case, cross-linking could be used to stabilize the interaction for subsequent immunoprecipitation. PflE could also act indirectly on FlgP, for example due to a signaling function or the recruitment of other factors required for basal disk assembly. One obvious candidate for further investigation would be FlgQ, which is essential for basal disk assembly via an unknown mechanism (Beeby et al., 2016; Sommerlad and Hendrixson, 2007), although it also does not seem to be a structural part of the flagellar disk complex (Drobnič et al., 2025). Besides the known requirement of FlgQ for FlgP stability and localization (Sommerlad and Hendrixson, 2007), the exact role of FlgQ remains unknown. However, our results indicate that PflE might act directly on FlgP, making the small protein a key component of the flagellar assembly pathway in *C. jejuni*.

Our work highlights not only the overall role of motility as an essential virulence trait for a major human pathogen, but also the role of a small protein in one of the most complex flagellar motor structures known to date. Given the elaborately complex system in *C. jejuni*, it serves as an ideal model organism to further explore small protein function in motility and virulence. The identification and characterization of PflE opens new avenues for understanding how small proteins can contribute to bacterial fitness and pathogenesis, a concept that extends well beyond *C. jejuni*.

## Materials and Methods

### Bacterial strains and culture conditions

*C. jejuni* strains (**Table S6**) were routinely grown either on Müller-Hinton (MH) agar plates or with shaking at 140 rpm in *Brucella* broth (BB) supplemented with 10 μg/ml vancomycin at 37°C in a microaerobic atmosphere (10% CO_2_, 5% O_2_). When required, MH agar was supplemented with the following antibiotics: 20 μg/ml chloramphenicol (Cm), 50 μg/ml kanamycin (Kan), 20 μg/ml gentamicin (Gm), or 250 μg/ml hygromycin B (Hyg). All *H. pylori* strains (**Table S6**) were grown on GC agar plates or in Brain Heart Infusion broth (BHI) liquid cultures with shaking at 140 rpm at 37°C in a microaerobic atmosphere (10% CO_2_, 5% O_2_). BHI was supplemented with 10 μg/ml vancomycin, 10% (v/v) FCS (fetal calf serum), 5 μg/ml trimethoprim, and 1 μg/ml nystatin. GC agar plates were supplemented with 10 μg/ml vancomycin, 10% (v/v) DHS (donor horse serum), 1 μg/ml nystatin, 5 μg/ml trimethoprim, 1% vitamin mix, and where appropriate, additional marker-selective antibiotics: 50 μg/ml kanamycin (Kan), or 20 μg/ml chloramphenicol (Cm). *E. coli* strains (**Table S7**) were grown aerobically at 37°C in lysogeny broth (LB) or on LB agar plates supplemented with the appropriate antibiotics for marker selection.

### Generation of a Tn5-based mutant pool in *C. jejuni* strain NCTC11168

A pool of transposon insertion mutants in *C. jejuni* strain NCTC11168 was generated as follows. First, a *Campylobacter* Tn5 transposon was constructed from the pMOD-2 plasmid (Epicentre) by replacing its native resistance cassette with the *aphA-3* kanamycin resistance cassette driven by the strong promoter of the *H. pylori* sRNA *repG* (Pernitzsch et al., 2014), and adding a T7 promoter. In detail, pMOD-2 was amplified by inverse PCR using CSO-1503 and CSO-1504 to introduce an outward-facing T7 promoter. After ligation and transformation into *E. coli* TOP10, the resulting plasmid (pSSv8) was verified by colony PCR with CSO-1500 and CSO-1504 and Sanger sequencing with PCRFWD and PCRREV. Next, the *aphA-3* kanamycin resistance cassette with the *H. pylori repG* promoter was amplified from pST1.1 (Dugar et al., 2018) using primers CSO-1567 and CSO-1568. Both pSSv8 and the resistance cassette insert were digested with *Eco*RI and *Hind*III, ligated, and transformed into *E. coli* to create plasmid pSSv9. Positive clones were verified by colony PCR with CSO-1500 and CSO-1501 and sequencing with PCRFWD and PCRREV. *In-vitro* transposition was then performed as follows. The *Campylobacter* Tn5 transposon, blunt-ended and 5’-phosphorylated, was amplified by PCR with Phusion polymerase with primers CSO-1500 and CSO-1501, each with 5’ phosphates. Genomic DNA was then purified from *C. jejuni* NCTC11168 wildtype with the Qiagen genomic-tip 100/G kit. Subsequently, *in-vitro* transposition was performed using EZ-Tn5 transposase (Epicentre) according to the manufacturer’s instructions with 0.25 pmol of transposon DNA and 1 µg of purified genomic DNA. The mutagenized DNA was purified by phenol:chloroform:isoamyl (25:24:1) alcohol extraction, and then insertion sites were repaired with T4 DNA polymerase (3 U), 10 mM dNTPs, and T4 DNA ligase (600 U) in T4 DNA ligase buffer (NEB). The repaired, mutagenized DNA was then used to transform *C. jejuni* NCTC11168 WT naturally as described previously (McLennan et al., 2008), and mutant colonies were recovered on MH agar plates containing kanamycin. The pool was harvested from plates with BB + vancomycin + 25 % (w/v) glycerol and stored at ‡80 °C. Colony PCR using CSO-1500 and CSO-1501 on genomic DNA from the pool verified the presence of the Tn5 transposable element. Genomic DNA was extracted from the pool grown once on MH plates with kanamycin and then subjected to library preparation and sequencing (see below) to identify and quantify insertion sites.

### Culture of eukaryotic cell lines

The Caco-2 human epithelial colorectal adenocarcinoma cells were routinely passaged in MEM medium (Gibco) with GlutaMAX™ and Earle’s salts, supplemented with 20% (w/v) fetal calf serum (FCS; Biochrom), 1% (w/v) Non-Essential Amino Acids (NEAA; Gibco), and 1% (w/v) Sodium Pyruvate (Gibco) in a 5% CO_2_ humidified atmosphere at 37°C. Cells were cultivated in T-75 (Sarstedt) flasks until 80 – 90 % confluence and passaged at a sub-cultivation ratio of 1/2 – 1/10. For passaging, cells were washed once with DPBS (without CaCl_2_ and MgCl_2_) to remove residual FCS and subsequently treated with 3 ml of 0.05% Trypsin-EDTA (Gibco) for 5 min at 37°C. The trypsinization process was stopped by addition of fresh cell culture medium to the flask and gentle mixing of the cells to give a homogenous single-cell suspension. Depending on the passaging ratio, a certain volume of this suspension was transferred to a new flask containing 10 – 13 ml of freshly supplemented cell culture medium and cells were given appropriate time to grow to confluence again.

### Generation of 3D tissue models

Caco-2 cell-based intestinal tissue models were generated as previously described (Alzheimer et al., 2020). Briefly, the decellularized extracellular matrix scaffold was fixed in cell crowns to create an apical and a basolateral compartment (SISmuc; (Linke et al., 2007; Schanz et al., 2010). Caco-2 cells were seeded in 500 µl cell culture medium onto the apical side of the scaffold, while the basolateral compartment was filled with 1.5 ml cell culture medium. Tissue models were then routinely cultivated either statically or dynamically on an orbital shaker (Celltron Infors HT) at 65 rpm in a 5% CO_2_ humidified atmosphere at 37 °C for 21 days. Fresh medium was supplied every two days.

### Infection of the 3D tissue model with *C. jejuni*

Tissue models were infected with *C. jejuni* wildtype and mutant strains as previously described (Alzheimer et al., 2020). Bacteria were grown as described above, harvested from liquid culture in mid-log phase (OD_600_ 0.4), and resuspended in cell culture medium to achieve an MOI of 20. From this bacterial cell suspension, serial dilutions were plated onto MH agar plates to determine the input amount of CFUs.

Bacterial suspensions were used to apically infect the tissue models and co-incubation was carried out in a 5% CO_2_ humidified atmosphere at 37°C without further mechanical stimulation. To isolate bacteria adherent to and internalized into the tissue models (ADH = adherence + internalization), infection was stopped at 24 hrs post infection and cell crowns were washed three times with DPBS to remove all non-adherent bacterial cells. Two tissue pieces per crown were collected using a tissue punch (Ø 5 mm, Kai Medical) and transferred to an Eppendorf tube with 500 µl of 0.1 % saponin in DPBS. The tissue pieces were incubated for 10 min at 37°C under agitation (1,500 rpm) to isolate bacteria from host cells. Serial dilutions were then plated on MH agar plates, colonies were counted, and CFU numbers were calculated as a percentage of input CFUs for each strain. To specifically isolate host cell-internalized *C. jejuni* (INT = internalization only), tissue models were additionally treated with 250 µg/ml gentamicin to kill extracellular bacteria for 2 hrs at 37 °C before CFUs were determined as described for ADH samples.

### Infection of 2D cell culture models with *C. jejuni*

2D Caco-2 cell monolayers were infected with *C. jejuni* wildtype and mutant strains as previously described (Alzheimer et al., 2020). Caco-2 cells were seeded into 6-well plates two days prior to the infection experiment. *C. jejuni* was grown in liquid culture to mid-log phase, resuspended in cell culture medium and used for infection of the epithelial monolayer at an MOI of 20. After infection, cells were washed three times with DPBS, lysed with 0.1 % saponin in 1 ml DPBS, and the resulting cell suspension was plated in serial dilutions on MH agar plates. For specific recovery of intracellular bacteria, cells were treated with 250 µg/ml gentamicin for 2 hrs at 37°C and CFUs were determined as described.

### Screen of the *C. jejuni* transposon mutant library in 3D and 2D infection models

The *C. jejuni* mutant pool was streaked onto MH agar plates from cryostocks and subsequently cultivated in BB liquid medium (incl. vancomycin and kanamycin) to mid-log phase (OD_600_ 0.4). This liquid culture was used to infect 12 Caco-2 tissue models and 2 x 6-well plates (*i.e.* 12 wells in total) with a confluent Caco-2 monolayer as described above for both model systems. After 4 hrs of infection in 2D and 24 hrs in 3D, the supernatant above the epithelial cells in each crown/well was harvested and pooled either for cell crowns (3D SUP) or 6-well plates (2D SUP). The bacteria in each supernatant pool were then plated on 12 x MH agar plates (145 x 20 mm) with vancomycin and kanamycin. The number of output CFUs recovered on each plate was approximated from a series of test infection experiments to result in approx. 4,000 colonies per plate. Next, bacterial CFUs both adherent to and internalized into the respective host cells (ADH) for half of the infected crowns and 6-well plates were isolated as described above for both 2D and 3D infection models. Bacterial cells harvested from these six tissue models or six wells were again pooled together and plated on 12 MH agar plates (145 x 20 mm) with vancomycin and kanamycin to recover 3D ADH and 2D ADH. After gentamicin treatment as described above, CFUs for internalized bacteria (3D/2D INT) were plated according to ADH samples. All CFUs were recovered for 48 hrs at 37 °C in a microaerobic environment and subsequently harvested, together with colonies from all plates representing an infection condition (2D or 3D: SUP, ADH, or INT) as follows. Next, 2 ml of fresh BB medium was pipetted onto each agar plate and colonies were carefully scraped off using an L-shaped Drigalski spatula (Hartenstein). Resulting bacterial suspensions for each condition were pooled in 50 ml Falcon tubes and well-mixed aliquots of approx. 10 OD_600_ were stored at ‡80 °C until subsequent extraction of gDNA and library preparation.

### Tn-seq library construction and sequencing

Genomic DNA was extracted from approximately 10 OD_600_ of thawed bacterial cells from each experimental output mutant pool (see description above) using the Qiagen Genomic-tip 100/G kit according to the manufacturer’s instructions. Purified DNA was resuspended in 1 x TE buffer and checked for quality by agarose gel electrophoresis. Next, 3 µg of purified DNA in 1 x TE was sheared by sonication in an ice bath to approx. 200 – 400 bp using a Bioruptor® Plus sonication device (Diagenode, Belgium) with low settings for 20 cycles of 30 sec on and 30 sec off. Following validation of shearing by electrophoresis on 6% PAA gels, the DNA was end-repaired and A-tailed using the NEBNext Ultra II DNA library Prep kit for Illumina® (New England Biolabs, E7645S) according to the manufacturer’s instructions. A Y-shaped splinkerette DNA adaptor (Devon et al., 1995), made by annealing CSO-3271 and CSO-3272 oligonucleotides (final concentration 15 µM), was then ligated to the sheared, repaired gDNA. Fragments of approximately 250 – 400 bp were then selected using SPRIselect beads (Beckman-Coulter, cat #B23317) according to the NEBNext Ultra II instructions (ratio of 0.4 x for the first selection and 0.2 x for the second selection) using a magnetic rack (Invitrogen). Transposon-chromosome junctions in size-selected, adaptor-ligated DNA (approx. 100 ng) were then amplified by PCR (19 – 22 cycles) with Phusion polymerase. Primers used for this enrichment PCR included a transposon-specific primer that adds the Illumina P7 sequence (CSO-2963), and a barcoding primer specific for the splinkerette adaptor (NEBNext Multiplex Oligos for Illumina®, Index Primer Sets 1 & 2) that adds a 6-nt long index and the Illumina P5 sequence. Following size selection from 6% PAA gels (250 – 500 bp), libraries were quantified using a Qubit v 2.0 fluorimeter and the HS dsDNA Assay Kit (ThermoFisher), as well as qPCR with the KAPA SYBR FAST Library Quantification Kit (KAPA Biosystems). Libraries were then sequenced on an Illumina® NextSeq 500 (high output, 75 cycles) at the Core Unit SysMed (University of Würzburg) with HPLC-purified custom primers (Sigma) for Read1 and Index sequencing (CSO-2964 and CSO-3305, respectively).

### Processing and mapping of Tn-seq reads

Raw Illumina reads in FASTQ format were trimmed from the 3‘ end with cutadapt (Martin, 2011) version 1.9.1/1.12 using a cut-off phred score of 20 (-q 20) to remove a combination of the splinkerette and Illumina adapter sequences (-a AGATCCCACTAGTGTAAGATCGGAAGAGCACACGTCTGAACTCCAGTCAC). After trimming, the READemption pipeline (Förstner et al., 2014) version 0.3.9/0.4.3 was applied to align all reads with a minimum length of 12 nt (-l 12) to the *C. jejuni* NCTC11168 genome (RefSeq Acc.-No: NC_002163.1) using the segemehl software version 0.2.0 (Hoffmann et al., 2009) with an accuracy cut-off of 95 % (-a 95). Subsequently, READemption was utilized to generate coverage plots representing the numbers of mapped reads per nucleotide, thereby considering only the 5’ position of uniquely aligned reads (-u -b first_base_only). Detailed information for read mapping statistics is given in **Table S8**.

### Definition of transposon insertion sites

Insertion of Tn5 results in a 9-bp target site duplication on both sides of the inserted transposon. As the custom primer used for sequencing can bind to either 3’ end of the palindromic transposon mosaic ends, read sequences of each insertion site should start with either 5’ end of the 9-bp direct repeat. This in turn causes a distance of 8 bp between read starts that map to different strands of the *Campylobacter* reference genome. *Bona fide* transposon insertion sites based on the genomic coverage plots were therefore defined as follows. For all positions n with coverage X_n_ on the forward strand, it was computationally checked if coverage Y_n+8_ on the reverse strand is > 0. If this was the case, an insertion site was annotated at the center position (n + 4) with a score of I_n+4_ = X_n_ + Y_n+8_. To normalize for sequencing depth, these scores were divided by the number of uniquely aligned reads for the respective library and multiplied by one million.

### Gene fitness evaluation based on Tn-seq-measured transposon insertions

The raw positional transposon insertion scores were converted to plot files using Artemis (http://www.sanger.ac.uk/resources/software/artemis/). This was necessary to achieve a compatible format for downstream analysis using the Bio-TraDIS package (Barquist et al., 2016). To map insertion sites to genes, NCBI and additional sRNA (Dugar et al., 2013) annotations were included in a Genbank file and, together with the insertion plot files, used as input for the tradis_gene_insert_sites analysis script, thereby omitting the last 10 % of mRNA coding regions (- trim3 0.1). Afterwards, the tradis_comparison.R script was used to compare gene fitness between pairs of growth conditions based on two biological replicates, requiring at least 10 read counts per gene and condition (-f -t 10).

### General recombinant DNA techniques and *C. jejuni*/*H. pylori* mutant construction

All plasmids generated and/or used in this study, as well as oligonucleotide primers (Sigma), are listed in **Tables S7** and **S9**, respectively. DNA constructs and mutations were confirmed by Sanger sequencing (Macrogen, Microsynth). Restriction enzymes, *Taq* polymerase for colony PCR (cPCR), and T4 DNA ligase were purchased from NEB. For cloning, Phusion high-fidelity DNA polymerase was used (Thermo Fisher Scientific). For *C. jejuni* and *H. pylori* mutant strain construction (deletion, chromosomal 3×FLAG-tagging, complementation by heterologous expression from the unrelated *rdxA* or the Cj0046 locus), double-crossover homologous recombination with DNA fragments introduced by electroporation or natural transformation was used. Resistance cassettes used for cloning were either *aphA-3* (Kan) (Skouloubris et al., 1998), *aph(7’’)* (Hyg) (Cameron and Gaynor, 2014), *C. coli cat* (Cm) (Boneca et al., 2008), or *aac(3)-IV* (Gm) (Bury-Moné et al., 2003). Details about *C. jejuni* and *H. pylori* mutant construction as well as transformation protocols are described below.

### Transformation of *C. jejuni* by electroporation

*C. jejuni* strains were grown from frozen stocks until passage one or two on MH agar plates containing the appropriate antibiotics. Cell material was resuspended in cold electroporation buffer (272 mM sucrose, 15% (w/v) glycerol), harvested by centrifugation (7000 x *g*, 5 min, 4°C) and washed twice. 50 μl of cell suspension were mixed with 500-1000 ng of purified PCR product. Cells were electroporated in a 1 mm-gap cuvette at 2.5 kV for 4-6 ms in a Bio-Rad MicroPulser and then immediately returned to ice. Cells were then transferred with BB media (supplemented with 10 μg/ml vancomycin) to non-selective MH plates, which were incubated overnight under microaerobic conditions at 37°C. After recovery, the cells were transferred to the appropriate selective MH agar plate and incubated until colony formation was observed. Obtained clones were validated by cPCR. Epitope tagging and complementation by heterologous expression strains were also verified by Sanger Sequencing (Microsynth, Macrogen).

### Natural transformation of *H. pylori*

Strains were grown from frozen stocks on GC agar plates, supplemented with appropriate antibiotics, until passage two. Afterwards the bacteria were streaked on a fresh GC agar plate (without additional selective antibiotics) in small circles and incubated for 6-8 hrs under microaerobic conditions and 37°C. Then, 500-1000 ng of PCR product was added to the cells on the plate and incubated overnight. To identify transformants, cells were transferred onto a selective GC plate and incubated until colonies were formed. gDNA was isolated and cPCR was used to validate positive clones. For complementation strains, clones were also validated by Sanger sequencing (Microsynth).

### Construction of *C. jejuni* and *H. pylori* deletion mutant strains

Deletion mutants of protein coding genes were constructed by replacement of most of the coding sequence with a non-polar resistance cassette by double-crossover homologous recombination. For this, a PCR product was generated via overlap PCR that fused a resistance cassette in between approx. 500 bp of sequence upstream and downstream of the target gene. Oligonucleotides used for deletion mutant construction are listed in **Table S9**. As an example, deletion of *flgP* (Cj1026c in *C. jejuni* strain NCTC11168 and HPG27_795 in *H. pylori* strain G27) with a non-polar Hyg cassette or non-polar Kan cassette is described.

The upstream region of Cj*flgP* was amplified with CSO-5130 and CSO-5131 from *C. jejuni* NCTC11168 WT genomic DNA (gDNA). The downstream region of Cj*flgP* was amplified with CSO-4236 and CSO-4237. The antisense primer of the upstream fragment and the sense primer of the downstream fragment introduced an overhang that overlaps with the resistance cassette at its 5’ and 3’ end, respectively. The Hyg resistance cassette was amplified from pACH1 (Cameron and Gaynor, 2014) with primers HPK1 and HPK2. Next, the three purified fragments (upstream, downstream, and resistance cassette) were fused by overlap PCR using primers CSO-5130 and CSO-4237 at a final concentration of 60 nM. For PCR reactions with the Hyg resistance cassette, 3% DMSO (dimethyl sulfoxide) was included. The purified PCR product was transformed into *C. jejuni* NCTC11168 WT strain (CSS-5295) by electroporation as described above. Putative *flgP*::*aph(7’’)* transformants (Δ*flgP*, CSS-6763) were verified by cPCR using CSO-5129 and CSO-2857. Deletion mutants in strain 81-176 (CSS-0063) were cloned in a similar manner.

The ∼500 bp upstream fragment of Hp*flgP* was amplified from *H. pylori* G27 WT gDNA using CSO-5998 and CSO-5999, while the downstream fragment was amplified using CSO-6000 and CSO-6001. The antisense primer of the upstream fragment and the sense primer of the downstream fragment introduced an overhang, which overlapped to the 5’ and 3’ end of the resistance cassette, respectively. The non-polar kanamycin resistance cassette (Kan) was amplified from pGG1 (Dugar et al., 2016) with HPK1 and HPK2. The three purified PCR products were fused together via overlap PCR using CSO-5998 and CSO-6001 as primers (final concentration of 60 nM). The obtained PCR fragment was purified and used for natural transformation into *H. pylori* G27 WT (CSS-0066) as described above. To verify the *flgP*::*aphA3* transformants (Δ*flgP*, CSS-8273), gDNA was isolated and used for cPCR with CSO-5997 and CSO-0023. All generated *C. jejuni* and *H. pylori* deletion strains are listed in **Table S6**. The non-polar Gm cassette was amplified using HPK1/HPK2 from pUC1813-apra (Bury-Moné et al., 2003). The non-polar Cm cassette was amplified using CSO-0613/CSO-0614 from pGD107.1 (Dugar et al., 2016).

### C-terminal 3xFLAG-tagging at the native locus

Epitope tagging at the native locus was performed via homologous recombination with an overlap PCR product. The PCR product included approximately 500 bp upstream of the open reading frame (ORF) penultimate codon fused to the 3×FLAG sequence and a Kan or Gm resistance cassette, followed by approximately 500 bp downstream of the ORF as described previously for Cj0978c-3xFLAG (PflE-3xFLAG, CSS-4716) (Froschauer et al., 2025) or Cj1643-3xFLAG (PflC-3xFLAG, CSS-4720) (Drobnič et al., 2025). Briefly, the construction of Cj0892c-3xFLAG (PflD-3xFLAG, CSS-4718) is described as an example. The upstream fragment (including the penultimate codon of Cj0892c) was amplified with CSO-3339/CSO-3771 from gDNA of *C. jejuni* NCTC11168 WT. The downstream fragment was amplified using CSO-3441/CSO-3442. The antisense oligo of the upstream fragment and the sense oligo of the downstream fragment added overhangs complementary to the 5’ end of 3xFLAG or to the 3’ end of the resistance cassette, respectively. A fusion of the 3xFLAG tag and the Kan resistance cassette was amplified from pGG1 (Dugar et al., 2016) using CSO-0065 and HPK2. Next, the three purified fragments (upstream, downstream, and 3x-FLAG with resistance cassette) were fused via a three-fragment overlap PCR. Overlap PCR was performed using primers CSO-3339 and CSO-3442 with a final concentration of 60 nM. Obtained clones were validated by cPCR using CSO-3343 and HPK2 and also verified by Sanger sequencing using CSO-0023 (Macrogen or Microsynth).

### Construction of *C. jejuni* and *H. pylori* complementation strains and point mutants

Complementation strains of *C. jejuni* were generated by heterologous expression of the gene of interest from the *rdxA* locus (Cj1066; (Ribardo et al., 2010)), if possible using the native promoter. Otherwise, the constitutive *metK* (Cj1096c) promoter was used. Constructs for complementation at *rdxA* were generated by insertion of the gene of interest into plasmids that already contained ∼500 bp of up- and downstream sequences complementary to the *rdxA* locus for homologous recombination (pSE59.1, pGD34.7) (Alzheimer et al., 2020; Dugar et al., 2018). The plasmid carried a chloramphenicol resistance cassette with its own promoter and terminator within the two *rdxA* homology regions. Oligonucleotides CSO-2276 and CSO-2277 were used to amplify constructs from generated plasmids for electroporation into *C. jejuni* and positive clones were validated by cPCR and Sanger sequencing. The construction of the Cj0892c (PflD) complementation plasmid pSSv106.5 as well as the Cj0892::*sfgfp* fusion (pSSv114.1) using the constitutive *metK* promoter were described previously (Drobnič et al., 2025). The complementation and sfGFP (Pédelacq et al., 2006) fusion constructs for Cj0978c (PflE, pSSv105.1 and pSSv113.1) and Cj1643 (PflC, pSSv107.8 and pSSv115.1) were cloned the same way. Additional examples for generation of *rdxA* plasmids and *C. jejuni* strains to drive expression with the native promoter at *rdxA* are described below.

Complementation with C-terminal 3xFLAG tagged Cj0978c (PflE) using its native promoter was achieved via generation of plasmid pKF5.3. The insert, including the *pflE* promoter region, its 5’UTR, as well as its coding sequence fused with a C-terminal 3xFLAG tag was amplified using CSO-5103 and CSO-5104 from a strain carrying the *pflE*-3xFLAG fusion at the native locus (CSS-4716). The plasmid backbone was amplified from pGD34.7 with CSO-3641 and CSO-0350 and *Dpn*I digested to remove template DNA. Both DNA fragments were digested with *Nde*I and *Xma*I, ligated, and transformed into *E. coli* TOP10. Clones were verified by cPCR (CSO-0065/3270) and Sanger sequencing (CSO-0643 or CSO-3270).

To generate point mutations in heterologously expressed genes, site-directed mutagenesis was performed by inverse PCR on *rdxA* plasmids with partially overlapping oligonucleotides that introduce the mutations. To generate pKF9.2 (expressing the PflE-M1 mutant; PflE-C17A-3xFLAG), inverse PCR using CSO-5123 and CSO-5124 was performed on pKF5.3. Purified DNA was *Dpn*I digested and directly transformed into *E. coli*. Introduction of the point mutation was validated by Sanger sequencing using CSO-3270. A similar strategy was used to generate the complementations for *flgP* (C-*flgP*, pKF14.1) and *flgP*-M1 (CjC17A, pKF16.1).

Several constructs for heterologous expression at *rdxA* were instead generated via overlap PCR, as described here for *flgQ*. The upstream fragment including *rdxA*, the Cm resistance cassette as well as the promoter and 5’UTR of *metK* was amplified with CSO-2276 and CSO-4207 from pSSv106.1. The downstream fragment including parts of *rdxA* was amplified from the same plasmid with CSO-1785/CSO-2277. The *flgQ* coding sequence was amplified with CSO-5171/CSO-5172 from *C. jejuni* NCTC11168 WT gDNA, which also added overhangs to the upstream and downstream fragments. Next, the three purified fragments were fused via a three-fragment overlap PCR using CSO-2276 and CSO-2277. Obtained clones were verified by cPCR (CSO-0644/CSO-0349) and Sanger sequencing (CSO-0644/CSO-3270).

Similarly, for complementations in *H. pylori*, a copy of the gene of interest was introduced in the *rdxA* locus (Goodwin et al., 1998) via double-crossover homologous recombination (Pernitzsch et al., 2014). Generation of the construct was performed via plasmid cloning in *E. coli*. Then, the region of interest was amplified from the plasmid with CSO-0017 and CSO-0018 and transformed via natural transformation into *H. pylori*. The description of the cloning strategy for complementation of *flgP* (HPG27_795; pKF53.3) is provided. The native promoter and coding sequence of *flgP* was amplified from *H. pylori* G27 WT gDNA using CSO-5173 and CSO-5174. The plasmid backbone was amplified from pSP109.6 (Pernitzsch et al., 2014) using CSO-2852 and CSO-6002. After purification of DNA fragments and *Dpn*I digestion of the plasmid template, insert and backbone were digested with *Nde*I and *Xma*I, ligated and transformed into *E. coli* TOP10 cells. Plasmid DNA was isolated and clones were verified by Sanger sequencing of the plasmid using CSO-0205.

To generate point mutations in heterologously expressed genes, the same strategy described above for *C. jejuni* was used. To generate the lipobox mutant of Hp*flgP* (Hp-C25A, pKF54.1), inverse PCR was performed using CSO-6003 and CSO-6004 and pKF53.3 as template DNA. Purified DNA was *Dpn*I digested and afterwards directly transformed into *E. coli* TOP10 cells. Plasmid DNA was isolated and sequenced using CSO-0205 to verify introduction of the point mutation.

### Cloning of transcriptional and translational reporter fusions for motility candidates

Transcriptional or translation reporter fusions to sfGFP (Pédelacq et al., 2006) were inserted into the Cj0046 pseudogene locus or *rdxA* locus, as described above for heterologous expression. The transcriptional sfGFP reporter fusion of P*_flgP_* fused to the unrelated 5’UTR of *metK* (pKF32.2) was cloned in an *rdxA*-based complementation plasmid (pSSv114.1, (Drobnič et al., 2025)) including the *sfgfp* gene and modified by overlap PCR for Cj0046 insertion. To generate pKF32.2, the promoter region of *flgP* (including the primary as well as a potential secondary *fliA* dependent promoter) was amplified with CSO-5604 and CSO-5605 from *C. jejuni* NCTC11168 WT gDNA, whereas the plasmid backbone was amplified using CSO-5096 and CSO-2989 from pSSv114.1. The backbone was first *Dpn*I digested and both fragments were subsequently digested with *Xma*I and ligated. The ligation product was transformed into *E. coli* TOP10 and clones were verified by cPCR (CSO-0644/3270) and sequencing (CSO-3270). For exchanging the flanking regions from *rdxA* to those of Cj0046, an overlap PCR was performed. The region containing the *flgP* promoter, *metK* 5’UTR and *sfgfp* was amplified from pKF32.2 using CSO-5606 and CSO-5100. The Cj0046 upstream region including a Gm^R^ cassette was amplified from pSSv54.3 (Svensson and Sharma, 2021) with CSO-1402/0484, while the downstream region of Cj0046 was amplified from the same plasmid using CSO-2751 and CSO-1405. All three fragments were fused by overlap PCR with CSO-1402/1405. The obtained *C. jejuni* clones were validated by cPCR (CSO-0833/3197) and Sanger sequencing (CSO-0833).

For generation of the *flgP* translational reporter (pKF34.1), the *flgP* 5’UTR as well as the first 10 codons of *flgP* were fused to *sfgfp*. The translational reporter was expressed from the unrelated *metK* promoter. Briefly, pKF34.1 was generated by inverse PCR on pSSv114.1 (Drobnič et al., 2025) using CSO-5684 and CSO-5685. By using these oligonucleotides that harbor overhangs, the 5’UTR of *metK* was exchanged with the one of *flgP* and additionally the first 10 codons of *flgP* were added to generate a translational fusion to sfGFP. The DNA fragment was *Dpn*I digested, ligated and transformed into *E. coli* TOP10. The obtained clones were validated by Sanger sequencing of plasmid DNA with CSO-0644. The fragment for transformation into *C. jejuni* was amplified using CSO-2276/2277. Obtained *C. jejuni* clones were validated by cPCR (CSO-0644/0349) and Sanger sequencing (CSO-3270).

### Construction of deletion mutants in *C. jejuni* strain 81-176

*C. jejuni* mutants were constructed by electroporation of plasmid DNA or natural transformation of *in vitro* methylated plasmid DNA following previously described methods (Beauchamp et al., 2017; Hendrixson et al., 2001). All plasmids were constructed by ligation of DNA fragments into plasmids by T4 DNA ligase or Gibson Assembly Mastermix (New England BioLabs).

pABT350 (Barrero-Tobon and Hendrixson, 2012) was electroporated into DRH2070/DRH2071 (81–176 *rpsL*^Sm^ Δ*flgP*/Δ*flgQ*) (Sommerlad and Hendrixson, 2007), resulting in DAR5947 (81–176 *rpsL*^Sm^ Δ*flgP vidA::cat-rpsL*) and DAR5950 (81–176 *rpsL*^Sm^ Δ*flgQ vidA::cat-rpsL*), respectively. A plasmid harboring an in-frame deletion of *pflE* was created by designing primers to amplify approximately 700 bases upstream and downstream of *pflE* and portions of *pflE* coding sequence to delete codon 2 to the penultimate codon. These DNA fragments were fused and assembled into *EcoR*I-digested pUC19 to create pDAR5909. pDAR5909 was then introduced into ABT366 (Barrero-Tobon and Hendrixson, 2012), DAR5947, and DAR5950. Transformants were isolated on MH agar with streptomycin and then screened for sensitivity to chloramphenicol. Colony PCR and DNA sequencing verified mutants that had replaced *vidA::cat-rpsL* with Δ*pflE* on the chromosome to result in DAR5971 (81–176 *rpsL*^Sm^ Δ*pflE*), DAR5969 (81– 176 *rpsL*^Sm^ Δ*pflE* Δ*flgP*), DAR6059 (81–176 *rpsL*^Sm^ Δ*pflE* Δ*flgQ*).

### Construction of PflE-1xFLAG plasmids for complementation studies in *C. jejuni* strain 81-176

Primers were designed to amplify by PCR from pRY109 (Yao et al., 1993) a 76-bp fragment containing the promoter for *cat* and its start codon, which was followed by an in-frame *Bam*HI restriction site and an in-frame 1xFLAG-tag epitope. This fragment was cloned into the *Xba*I and *Pst*I sites of pRY112 (Yao et al., 1993) to result in pDAR5476 to allow for expression of C-terminal 1xFLAG-tagged proteins for complementation. Primers were designed to amplify the coding sequence of *pflE* from codon 2 through the penultimate codon with both 5′ and 3′ in-frame *Bam*HI sites from the *C. jejuni* 81–176 genome. The resultant DNA fragment was cloned into the *Bam*HI-digested pDAR5476 to create pDAR5951.

pDAR5476 and pDAR5951 were transformed into *E. coli* DH5α/pRK212.1, which served as the donor strain for conjugation into *C. jejuni* (Figurski and Helinski, 1979). Plasmids were then conjugated into *C. jejuni* strains DAR5971, DAR5969 and DAR6059 as previously described (Guerry et al., 1994), resulting in paired vector and PflE-1xFLAG expressing strains in Δ*pflE* (DAR6023 and DAR6029), Δ*pflE* Δ*flgP* (DAR6013 and DAR6019), and Δ*pflE* Δ*flgQ* (DAR6067 and DAR6071).

### Total protein analysis by SDS-PAGE and western blotting

Bacterial cells were grown to mid-exponential phase (for motile strains OD_600_ of ∼0.4, for non-motile strains OD_600_ ∼0.6), unless otherwise stated, and harvested by centrifugation (13,000 rpm, 5 min, 4°C). Cell pellets were resuspended in 1× protein loading buffer (PL; 62.5 mM Tris-HCl, pH 6.8, 100 mM DTT, 10% (v/v) glycerol, 2% (w/v) SDS, 0.01% (w/v) bromophenol blue). Samples corresponding to an OD_600_ of 0.1 were separated on 12% SDS-polyacrylamide (PAA) gels and blotted to a nitrocellulose membrane (semidry system, Peqlab). After blocking with 10% (w/v) milk powder in TBS-T (Tris-buffered saline-Tween-20), membranes were incubated overnight with primary antibody in 3% BSA/TBS-T [monoclonal mouse anti-FLAG (1:1,000; Sigma-Aldrich, #F1804-1MG), monoclonal mouse anti-GFP (1:1,000, Roche #11814460001), anti-GroEL (loading control, rabbit polyclonal, 1:10,000, GE Healthcare, #RPN430) or FlgP antiserum (rabbit, 1:20,000, (Sommerlad and Hendrixson, 2007))] at 4°C. As secondary antibody either anti-mouse IgG horseradish peroxidase (HRP) conjugate (sheep polyclonal, 1:10,000; GE Healthcare, #RPN4201) or anti-rabbit IgG HRP conjugate (goat polyclonal, 1:10,000; GE Healthcare, #RPN4301) in 3% BSA/TBS-T was used. Blots were developed using enhanced chemiluminescence reagent on a LAS-4000 Chemiluminescence Imager (GE Healthcare). For size estimation, a Prestained Protein Marker (Thermo Fisher Scientific) was loaded.

### RNA isolation and northern blotting

Total RNA was extracted using the hot phenol method as previously described (Sharma et al., 2010). Briefly, *C. jejuni* cultures were grown to log phase in Brucella broth and culture corresponding to approximately 4 OD_600_ was mixed with 0.2 volumes of stop-mix (95% ethanol and 5% phenol, v/v). After snap-freezing in liquid nitrogen, samples were stored at −80°C until RNA extraction. Frozen samples were thawed on ice and centrifuged at 4°C (4,500 x *g*, 20 min). Cell pellets were then lysed by resuspension in 600 μl of a solution containing 0.5 mg/ml lysozyme and 1% SDS in Tris-EDTA buffer (pH 8.0) and incubation for 2 min at 64°C. Total RNA was extracted from the lysate using the hot-phenol method (Sharma et al., 2010). For northern blot analysis, total RNA was treated with DNase I (Thermo Fisher Scientific) to remove residual genomic DNA, and 5–10 μg of digested RNA in Gel Loading Buffer II (GLII, Ambion) was loaded on 6% polyacrylamide (PAA)/7 M urea denaturing gels in 1× TBE buffer. Following electrophoretic separation, RNA was transferred to Hybond-XL membranes (GE Healthcare) by electroblotting and cross-linked to the membrane via UV light. The membrane was hybridized with γ^32^P-ATP end-labeled DNA oligonucleotides (**Table S9**) in Roti Hybri-quick (Roth) at 42°C overnight. Membranes were then washed at 42°C with decreasing concentrations of SSC (saline-sodium citrate) + 0.1% SDS buffer, dried, and exposed to a PhosphorImager screen. Screens were scanned using a FLA-3000 Series PhosphorImager (Fuji) and bands were quantified using AIDA software (Raytest, Germany).

### Motility assays

For motility assays of *C. jejuni*, strains were grown in Brucella broth (BB) with shaking to mid-log phase at 37°C in a microaerobic environment. Soft-agar plates (BB + 0.4% Difco agar) were inoculated with 1 µl of bacterial culture and incubated right-side-up for approx. 24 hrs post-inoculation. The swimming halo radius of each strain was measured two times (perpendicularly) and averaged to provide measurements of the technical replicates. In general, each motility assay was performed at least in two biological replicates (from independent cultures). Statistical analysis was performed with GraphPad Prism software using an unpaired, two tailed Student’s *t*-test to assess significance. *C. jejuni* WT strain and a *flaA* deletion mutant were always included as motile and non-motile controls, respectively.

Motility assays for *H. pylori* were performed similarly. Briefly, *H. pylori* strains were grown to log phase in BHI liquid cultures as described above and inoculated in soft-agar plates (BB broth + 0.4% Difco agar, 10 μg/ml vancomycin, 5% (v/v) FBS, 5 μg/ml trimethoprim, 1 μg/ml nystatin). Then, 0.5 OD_600_ of log-grown bacterial cultures were harvested by centrifugation (6,500 x *g*, 10 min at room temperature) and resuspended in 1 ml supplemented BHI. 1 µl of the OD_600_ adjusted suspension was used to inoculate the soft-agar plates. Motility assays were performed in three biological replicates (independent cultures) as well as in three technical replicates. The plates were incubated for approximately one week (5-7 days) right-side-up at 37°C under microaerobic conditions. Measurements, statistics and controls were done as described for *C. jejuni*.

### Transmission electron microscopy (TEM)

*C. jejuni* strains grown on MH agar plates with the appropriate antibiotics were very gently resuspended in 1 ml DPBS (Dulbecco’s phosphate buffered saline) and harvested by centrifugation at 5,000 x *g* for 5 min at RT. The cell pellet was subsequently resuspended in 2% glutaraldehyde in 0.1 M cacodylate buffer and incubated overnight at 4°C. The following day, a 1:10 bacterial cell suspension was mounted onto hydrophilized TEM grids. After a washing step with H_2_O for 1 min, *C. jejuni* cells were stained with 2% uranyl acetate and finally inspected using a Zeiss EM10 transmission electron microscope.

### Subcellular localization analysis of sfGFP-fusion proteins by microscopy

Superfolder GFP fusion complementation strains of Cj0892c, Cj0978c, and Cj1643 were grown under agitation at 140 rpm in Brucella Broth liquid culture in a microaerobic environment until mid-log phase (OD_600_ of 0.4). Bacterial cells were then harvested by centrifugation (6,500 x *g*, 5 min, RT), washed once with DPBS, and fixed with 2% PFA for 2 hrs at RT in the dark. After three washing steps with DPBS, microscopy images were taken using a Leica SP5 confocal laser scanning microscope.

### Cell fractionation of *C. jejuni*

The subcellular fractionation protocol is based on two previously published protocols for *C. jejuni* (Bingham-Ramos and Hendrixson, 2008; Sommerlad and Hendrixson, 2007) and allowed separation of the cytoplasm, the periplasm, the inner membrane, and the outer membrane. As control for successful fractionation, different antisera against known localised proteins were probed. Here, outer membrane protein FlgH (anti-FlgH; UTGP161), periplasmic protein Cjj81176_0382 (anti-Cjj81176_0382; M17, (Bingham-Ramos and Hendrixson, 2008)), inner membrane protein FliF (anti-FliF; M202, (Boll and Hendrixson, 2013)) and cytosolic protein RpoA (anti-RpoA; UTGP275, (Waller et al., 2024)) or polyclonal GroEL antibody were used. A 16 as well as a 4 OD_600_ cell pellet was harvested by centrifugation (4°C, 5,000 rpm, 20 min) from standard cultures grown to exponential phase (OD_600_ 0.3-0.4) for preparation of the periplasmic and cytoplasmic fractions, as well as the inner and outer membrane fraction, respectively. In addition, a whole cell lysate control, where cells are boiled in 1 x PL, was harvested. The 4 OD_600_ pellet for the membrane preparation was washed once with 1 ml 10 mM HEPES, pH 7.4 and stored at ‡20°C until further usage. The 16 OD_600_ pellet for periplasmic and cytoplasmic fractions required immediate processing and was washed twice in 1 ml PBSG (PBS with 0.1% gelatin). Afterwards, the cell pellet was resuspended in 400 μl PBSG including 2 mg/ml polymyxin B sulfate (Sigma), and was incubated rotating for 30 min at 4°C. After centrifugation (30 min, 15,000 rpm, 4°C) the supernatant containing the periplasmic proteins, was recovered. An additional centrifugation step (30 min, 15,000 rpm, 4°C) of the periplasmic fraction was performed to avoid contamination with spheroplasts. The spheroplast-containing pellet was washed with 500 μl 10 mM HEPES pH, 7.4, resuspended in 1 ml 10 mM HEPES, pH 7.4, and lysed by sonication. For sonication, 4 x 30 s cycles were used and after each cycle a break of 30 s followed. During the whole sonication process, the sample was constantly cooled in an ice water slurry. Settings on the sonicator were as follows: 50% cycle, 50% power. The lysate was centrifuged to remove unbroken cells (2 min, 15,000 rpm, 4°C). The obtained supernatant was centrifuged two more times to remove the membrane particles. The first centrifugation step was for 1 h (15,000 rpm, 4°C), whereas the second centrifugation was 30 min (15,000 rpm, 4°C) to allow pelletting of membranes. The cytoplasmic fraction was obtained from the supernatant of the second centrifugation step.

The 4 OD_600_ cell pellet was thawed on ice, resuspended in 1 ml 10 mM HEPES pH 7.4 and lysed by sonication (see above). The obtained lysate was centrifuged to remove unbroken cells (2 min, 15,000 rpm, 4°C) and further centrifuged (1 h, 15,000 rpm, 4°C) to pellet the membrane. The membrane pellet was washed with 500 μl 10 mM HEPES pH 7.4. To separate the inner and the outer membrane proteins, the membrane pellet was resuspended in 200 μl 1% N-lauroylsarcosine sodium salt in 10 mM HEPES pH 7.4 and was incubated slowly shaking for 30 min at room temperature to solubilize inner membrane proteins. The outer membrane was harvested by centrifugation (1 h, 15,000 rpm, 4°C). To avoid cross-contamination, the supernatant, containing inner membrane proteins, was centrifuged again (30 min, 15,000 rpm, 4°C) to remove potential left over outer membrane particles. The supernatant was saved as the inner membrane fraction. The outer membrane pellet was washed once with 500 μl 10 mM HEPES and was saved as outer membrane fraction. For analysis, an aliquot of each fraction representing 0.32 OD_600_ of cells was loaded on a western blot. For the whole cell lysate, 0.16 OD_600_ was loaded.

### FlgH purification

The coding sequence of *C. jejuni* 81-176 *flgH* from codon 20 to the penultimate codon was amplified by PCR from *C. jejuni* 81–176 and then cloned into *Bam*HI/*Xho*I-digested pET21a (Novagen) to create pDAR2575. This plasmid was then moved to *E. coli* BL21(DE3) for recombinant FlgH-6xHis purification. BL21(DE3) containing pDAR2575 was grown to mid-log phase in LB broth with shaking at 37°C and then induced with 1-mM isopropyl-beta-D-1-thiogalactopyranoside (IPTG) for 3 h at 37°C. After induction of protein, cell pellets were collected by centrifugation and resuspended in 8M urea. Cells were passed through an EmulsiFlex-C5 disruptor at 15,000–20,000 lb/in^2^ to lyse cells. The soluble fraction was recovered by centrifugation at 20,000 × *g* for 30 min. Recombinant FlgH was purified with Ni^2+^-agarose beads after washing with 8M urea, pH 6.3 and elution of FlgH-6XHis with 8M urea, pH 5.5. Glycerol was added to a final concentration of 20% prior to storage at −80°C.

### FlgH polyclonal antisera generation

Purified recombinant FlgH-6xHis was used to immunize guinea pigs by standard procedures for antisera generation via a commercial vendor (Cocalico Biologicals), resulting in antisera UTGP161. All uses of animals in experimentation were approved by The Institutional Animal Care and Use Committee (IACUC) at the University of Texas Southwestern Medical Center.

### Protein stability assays

Protein stability assays were modified from previously published protocols (Geuskens et al., 1992; Griffith et al., 2004). *C. jejuni* strains were grown to an OD_600_ of 0.4 or 0.6 (exponential phase) for motile and non-motile strains, respectively. Then, chloramphenicol (Cm) or spectinomycin (Spc) was added to a final concentration of 100 μg/ml and time point zero was immediately harvested (2 ml of culture) by centrifugation (13,000 rpm, 5 min, 4°C). The supernatant was discarded and the cell pellet was resuspended in 1 x PL to obtain 0.01 OD_600_/μl. Samples were boiled at 95°C for 8 min shaking at 1000 rpm. Additional time points were collected after 10, 30, 60, 120, 180 min after addition of Cm or Spc, respectively.

### Rifampicin assays

To determine transcript stability *in vivo*, strains were grown to an OD_600_ of 0.5 (mid-log phase) and treated with rifampicin to a final concentration of 500 μg/ml. Samples were harvested for RNA isolation as described above at indicated time points following rifampicin addition (0, 2, 4, 8, 16, 32, 64 min). Time point zero was immediately harvested after addition of rifampicin. After isolation, residual DNA was removed from RNA by treatment with DNase I (Thermo Fisher Scientific) according to the manufacturer’s recommendations. Five micrograms of each DNase I-digested RNA sample was used for northern blot analysis as detailed above. Rifampicin assays were performed in biological duplicates.

### Bacterial culture and cryoET sample preparation

All *C. jejuni* NCTC11168 mutants used for cryoET were grown from frozen stocks on MH agar supplemented with trimethoprim (10 µg/mL) for 1-2 days at 37°C in microaerophilic conditions (5% O_2_, 10% CO_2_, 85% N_2_). Cells were re-streaked on fresh plates and grown for an additional 16 h. Fresh cells were suspended from MH plates into ∼1.5 mL of MH broth and concentrated to an approximate theoretical OD_600_ of 10 by centrifugation at 3000 rpm for 5 min on a tabletop microcentrifuge and resuspending appropriately. 30 µL of the concentrated cell sample was mixed with 10 nm gold fiducial beads coated with BSA. 3 µL of this mixture was applied to freshly glow-discharged R2/2, 300 mesh Quantifoil grids. Grids were plunge-frozen in liquified ethane-propane using a Vitrobot mark IV.

### cryoET data collection

Tilt series were collected on a 200-keV F20 TEM microscope equipped with a Falcon II detector (Thermo Fisher, formerly FEI) and either an Elsa or 626 side-entry cryo-holders (Gatan). Data collection was automated with Leginon software (Carragher et al., 2000), implementing a bidirectional tilt scheme from ‡57_o_ to + 57° in 3° increments, starting at +18°. Images were collected at 29,000x magnification, giving a pixel size of 7.002 Å.

### Tomogram reconstruction and subtomogram averaging

Automated tomogram reconstruction was implemented with in-house custom scripts, using IMOD (Kremer et al., 1996) and Tomo3D (Agulleiro and Fernandez, 2015). Low-pass filtering to 3.5 nm was applied to all data before reconstructing tomograms with the SIRT method. For all three structures, subtomogram averaging was performed using the Dynamo package (Castaño-Díez et al., 2012). Motors were manually annotated as ‘oriented particles’ and extracted for averaging. All alignment and averaging steps were performed on un-binned tomograms. For each structure, an initial model was obtained by reference-free averaging of the oriented particles, with randomized z-axis rotation to alleviate missing wedge artifacts. This initial model was used for a first round of alignment and averaging steps, implementing an angular search and translational shifts, with cone diameter and shift limits becoming more stringent across iterations. The resulting average was used as a starting model for a round of finer alignment and averaging. In this round, custom alignment masks were implemented, focusing on the periplasmic and inner membrane-associated parts of the motor. This excluded dominant features that would otherwise drive the alignment, most prominently the outer membrane. 17-fold rotational averaging was applied, as it is established that the *C. jejuni* motor displays 17-fold rotational symmetry (Beeby et al., 2016). The final averages are derived from 195 particles for Δ*pflD*, 96 particles for Δ*pflE*, and 101 particles for Δ*pflC*.

### Motility protein homolog identification and sequence analysis

Homologs of motility proteins were mainly identified by BLASTp against the NCBI Refseq protein database using the Refseq non-redundant database with less stringent parameters (E-value <10, word size 2) using taxIDs in **Table S4**. Homologs with coverage of at least 50% were retained. For species missing homologs at this cutoff, we used manual inspection based on BLASTp with a homolog from a more closely related taxon, synteny, KEGG orthology (https://www.genome.jp/kegg/) (Kanehisa and Goto, 2000), or previously published investigations (Chaban et al., 2018; Mo et al., 2022). Some *Helicobacter* homologs of Cj0892c were identified by BLASTp with the putative *H. pylori* homolog HP0398, which shows similar synteny to Cj0892c. Cj0978c matches >100 aa and other protein matches of extremely long comparative length were discarded. Sequences were aligned using MultAlin (http://multalin.toulouse.inra.fr/multalin/) (Corpet, 1988). To generate a phylogenetic tree based on FlgP homologs, the amino acid sequence of *Cj1026c* from *C. jejuni* NCTC11168 was used to search the KEGG database (https://www.genome.jp/tools/blast/, (Kanehisa and Goto, 2000)) using BLASTp using default parameters. Species shown in **Figure 5A** were selected and used to build a phylogenetic tree using the KEGG workflow mafft_default-none-none-fasttree_default as described. The tree was visualized with iTOL (Letunic and Bork, 2024). KEGG species tags are listed in **Table S4**.

## Acknowledgements

We thank Lars Barquist for input into Tn-seq data analysis, members of the Nicolai Siegel lab for assistance with library preparation, and Claudia Gehrig, Imaging Core Facility, Biocenter, University of Würzburg for TEM assistance. Vivian Monzon and Ann-Janine Imsieke generated several plasmids for this study.

## Author Contributions

K.F., S.L.S., M.A., and C.M.S. designed the research. K.F., S.L.S, M.A., D.A.R., E.H., L.H., and T.D. performed lab work. K.F., S.L.S., M.A., T.B., T.D., and L.H. analyzed data. K.F., S.L.S., M.A., and C.M.S. interpreted the data and wrote the manuscript, which comments from co-authors. D.R.H. provided resources, M.B., and C.M.S. supervised research and provided resources.

