## Supplementary Figures for "A small protein essential for flagellar assembly and virulence of *Campylobacter jejuni*"

**This document contains:**

**Figures S1-S10**

**Supplementary References**

### Supplementary Figures

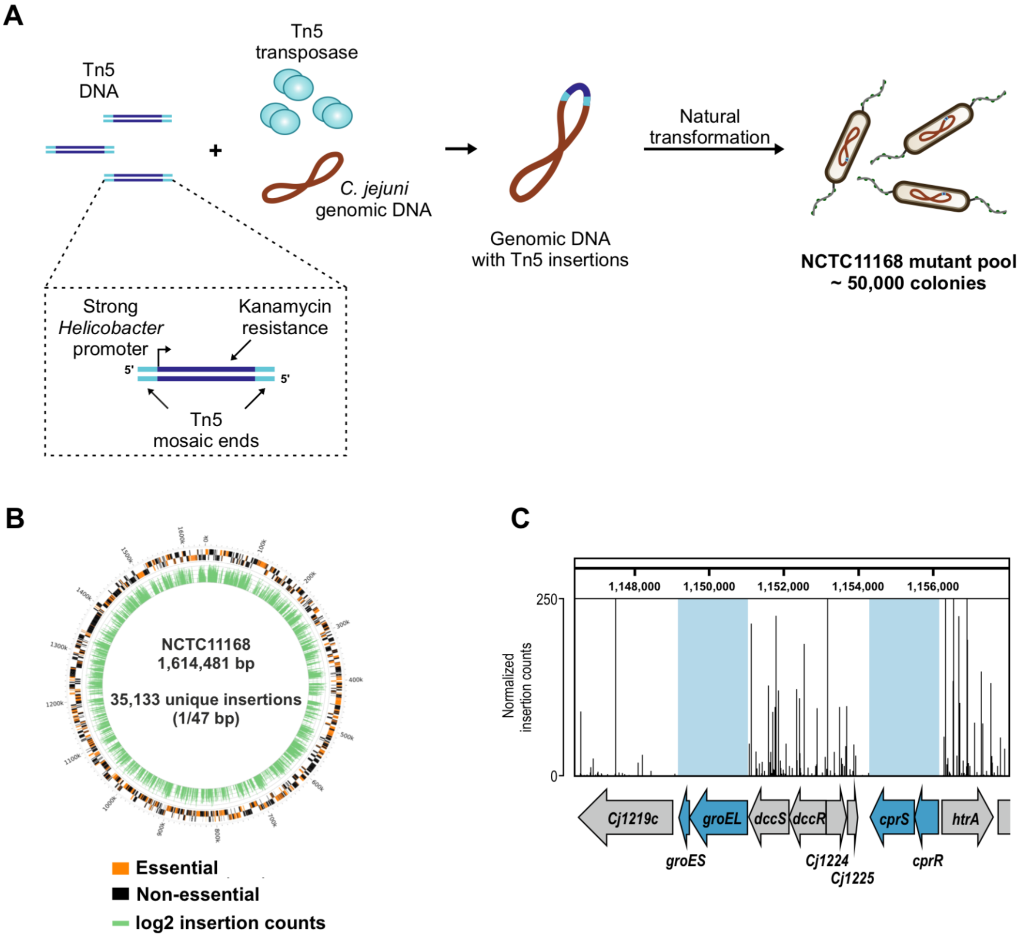


**Figure S1. Design and validation of a *C. jejuni* strain NCTC11168 Tn5 mutant pool. (A)** Schematic of Tn5-based transposon mutant library generation by *in-vitro* transposition into genomic DNA (gDNA) from the *C. jejuni* wild-type strain NCTC11168. A Tn5 transposon was modified to include a kanamycin resistance cassette (*aphA-3*) under control of the strong *repG* promoter from *Helicobacter pylori* (Pernitzsch et al., 2014). A transcriptional terminator was not included. *In vitro*-mutagenized gDNA was naturally transformed into *C. jejuni* NCTC11168 and ~50,000 kanamycin-resistant colonies were collected. For more details, see **Methods**. **(B)** Tn insertion site counts per gene (green) in the *C. jejuni* NCTC11168 chromosome (NC_002163.1) as well as genes called as essential/non-essential (orange/black) for the *C. jejuni* Tn5 mutant pool. The plot was generated using Circos (Krzywinski et al., 2009). The height of the green bar indicates the log2 of the number of insertions at that gene (insertion density). Essential genes have no insertions. bp: basepairs. **(c)** Read coverage screenshots for loci harboring previously described essential genes, representing insertion counts from the Tn-seq analysis of the mutant library mapped to its reference genome. Blue arrows: putative essential genes *groES*, *groEL*, *cprS*, and *cprR* based on our study. While *groES*, *groEL*, and *cprR* were identified as essential in at least one other study (de Vries et al., 2017; Gao et al., 2014; Mandal et al., 2017; Metris et al., 2011; Stahl and Stintzi, 2011), essentiality of *cprS* in strain NCTC11168 was unique to our dataset, but was found to be essential in strain 81-176 in two other studies as well (de Vries et al., 2017; Mandal et al., 2017). Coverage was visualized with Integrated Genome Browser (IGB) (Freese et al., 2016).

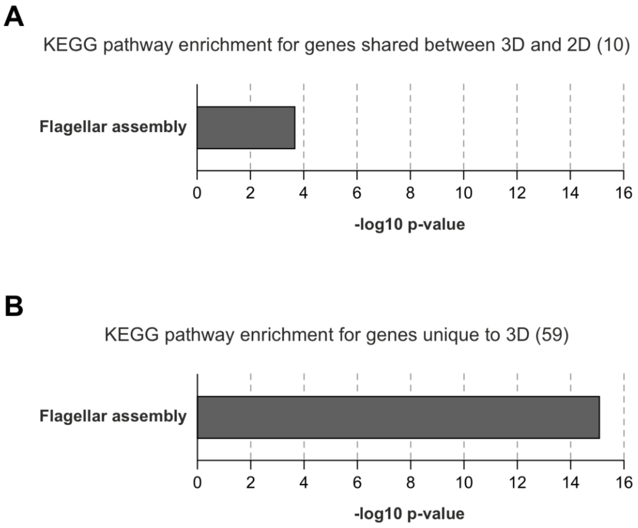


**Figure S2. Functional enrichment analysis for genes with decreased fitness identified in the Tn-seq screen. (A, B)** Functional enrichment for KEGG pathways was analysed using STRINGdb (Szklarczyk et al., 2023) on all genes, whose disruption by Tn5 insertion resulted in decreased fitness (ADH+INT) in either both 3D and 3D infection model **(A)** or in the 3D tissue model only **(B)**. Decreased fitness is defined as a significant ($p \leq 0.05$) log2FC ≤ -1 of gene-wise transposon insertion counts obtained from libraries prepared from adherent and internalized samples (ADH) or internalized samples only (INT) versus those from the supernatant bacteria (SUP; ≥ 10 reads). KEGG pathways with a Benjamini-Hochberg adjusted p-value ≤ 0.05 are shown and depicted as -log10 of their p-value.

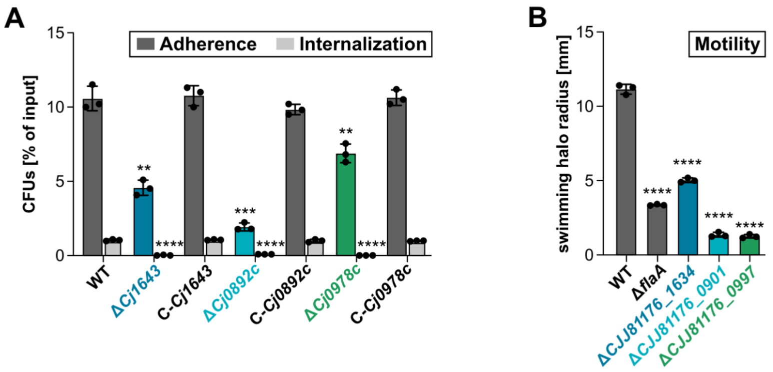


**Figure S3. Infection assays in 2D Caco-2 monolayer and motility-phenotypes of Tn-seq candidates in *C. jejuni* strain 81-176. (A)** Validation of Tn-seq infection phenotypes for *Cj1643*, *Cj0892c*, and *Cj0978c* mutants in 2D Caco-2 monolayers. CFUs were isolated for adherent and internalized bacteria (Adherence) or internalized bacteria only (Internalization) of NCTC11168 wildtype (WT), deletion (Δ), and complementation (C) strains after 4 hrs p. i. CFUs are displayed as a percentage of input bacteria and represent the mean of three biological replicates with corresponding SDs. Significance is given with respect to the parental WT. **(B)** Motility assays in soft agar for *C. jejuni* strain 81-176 wildtype (WT) and deletion mutants of *CJJ81176_1634* (*Cj1643*), *CJJ81176_0901* (*Cj0892c*), and *CJJ81176_0997* (*Cj0978c*). Δ*flaA*: non-motile control. Mean of three biological replicates and SD are shown. ****: $p < 0.0001$, ***: $p < 0.001$, **: $p < 0.01$ (Student's *t*-test).

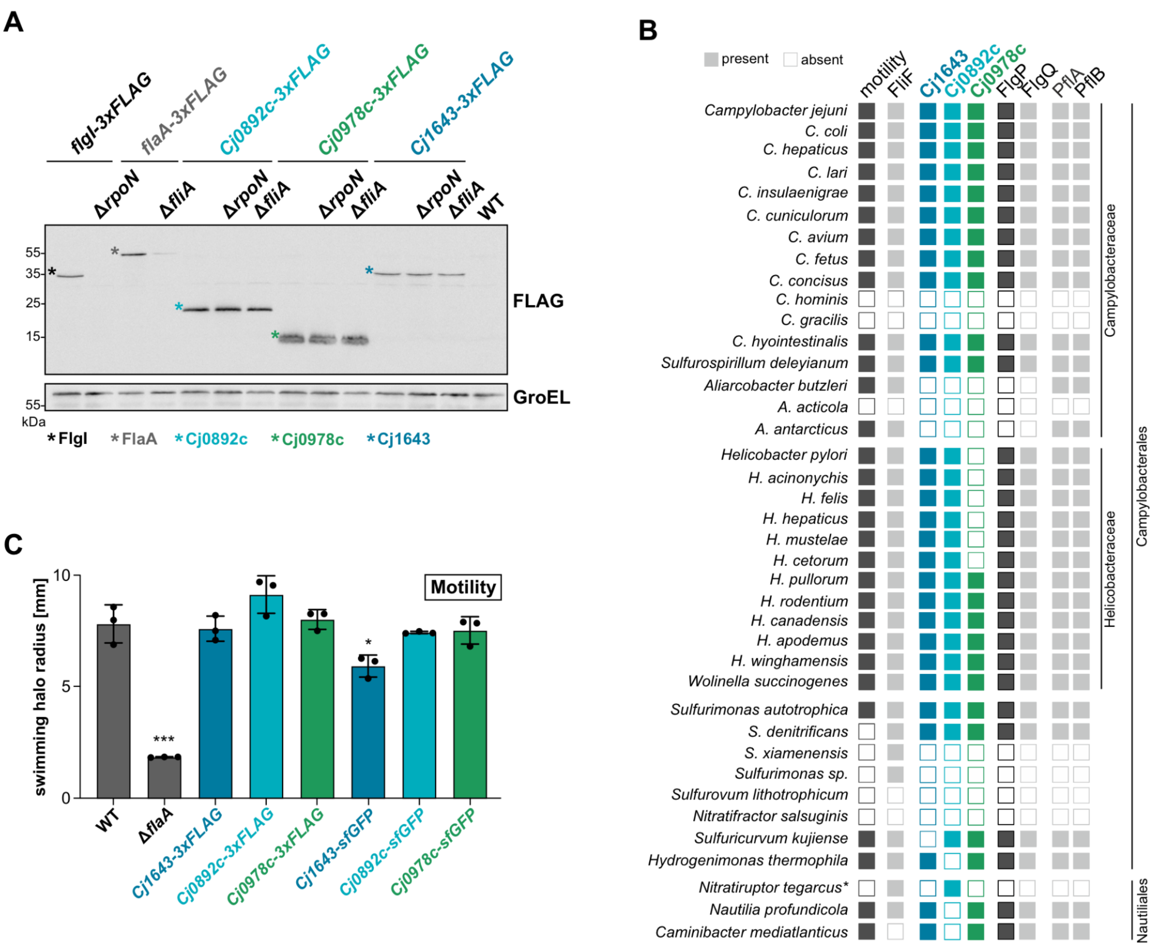


**Figure S4. Examination of flagella co-regulation, conservation of motility-associated Tn-seq candidates, and motility. (A)** Western blot analysis of Cj0892c-3xFLAG, Cj0978c-3xFLAG, Cj1643-3xFLAG, FlgI-3xFLAG (RpoN-dependent control), and FlaA-3xFLAG (FliA-dependent control) in WT as well as Δ*rpoN* and Δ*fliA* flagella sigma factor deletion backgrounds. Representative of two independent experiments. FLAG-tagged proteins were detected with an anti-FLAG antibody. GroEL served as a loading control. WT: untagged control. **(B)** Conservation analysis of Cj1643, Cj0892c, Cj0978c, and relevant flagellar proteins in Campylobacterota. Filled squares indicate the presence of respective genes in the indicated species as well as whether they are motile. Asterisk: the strain analyzed is non-motile due to a transposon insertion in a flagellin gene (Sievert et al., 2008). See **Methods** and **Table S4** for details. **(C)** Motility assays in soft agar for *C. jejuni* NCTC11168 WT and 3xFLAG/sfGFP fusions of *Cj0892c*, *Cj0978c*, and *Cj1643*. Tags were fused to the respective C-terminus of the genes and either integrated at the natives locus (for 3xFLAG) or expressed as complementation constructs (for sfGFP) from the unrelated *rdxA* locus in their respective deletion backgrounds. Δ*flaA*: non-motile control. Depicted are the mean and SD of three biological replicates. Significance vs. WT, ***: $p < 0.001$, *: $p < 0.05$ (Student's *t*-test).

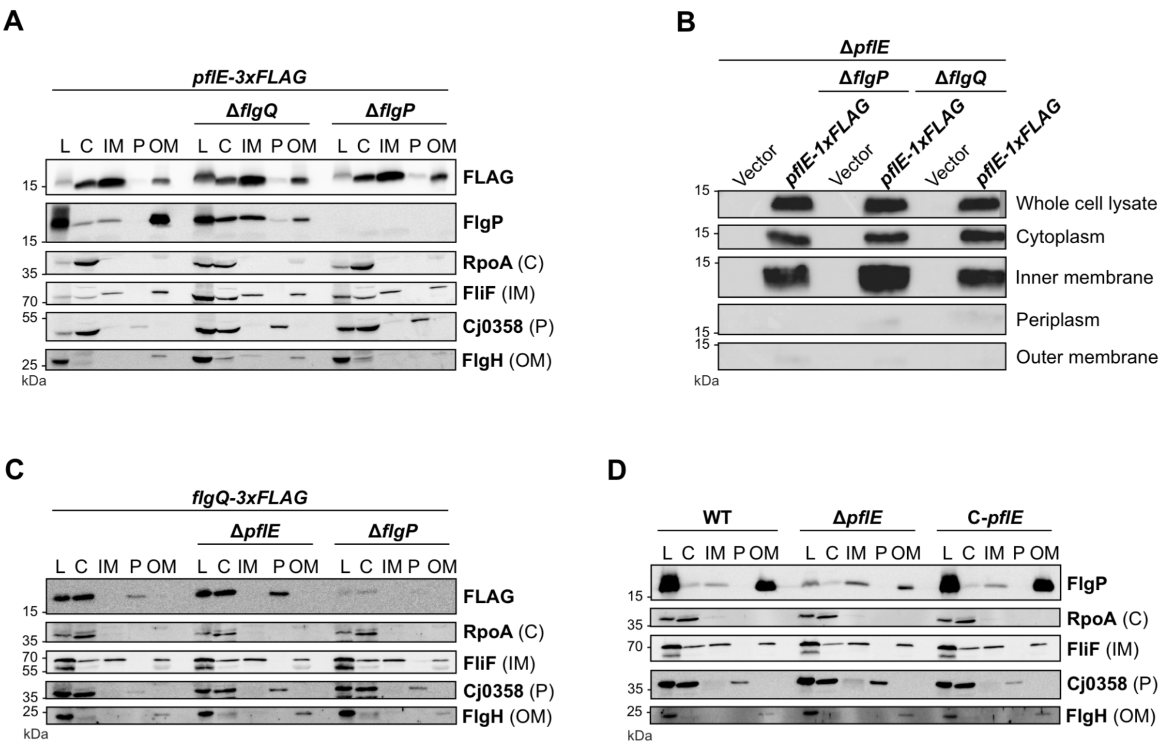


**Figure S5. Subcellular localization of PflE, FlgQ, and FlgP. (A, C-D)** Subcellular localization of PflE-3xFLAG **(A),** FlgQ-3xF **(C)**, and FlgP **(D)** in NCTC11168 wildtype, *flgQ*/*flgP*/*pflE* deletion (Δ), or *pflE* complementation (C-*pflE*) backgrounds. Strains were subjected to subcellular fractionation and analyzed by western blot. FlgP was detected with a specific antiserum while PflE and FlgQ were detected with anti-FLAG. Controls for fractionation: (C) cytosolic protein RpoA (anti-RpoA; UTGP275; (Waller et al., 2024)), (IM) inner membrane protein FliF (anti-FliF; M202; (Boll and Hendrixson, 2013)), (P) periplasmic protein Cj0358 (anti-CJJ81176_0382; M17; (Bingham-Ramos and Hendrixson, 2008)), and (OM) outer membrane protein FlgH (anti-FlgH; UTGP161). **(B)** Western blot analysis of fractionation samples of PflE-1xFLAG in *C. jejuni* strain 81-176. PflE-1xFLAG was expressed from a plasmid under control of a constitutive promoter for *cat* (encoding chloramphenicol acetyltransferase) in Δ*pflE*, Δ*flgP* Δ*pflE*, and Δ*flgQ* Δ*pflE* deletion mutant backgrounds. Vector refers to the empty plasmid control expressed in the same strain backgrounds.

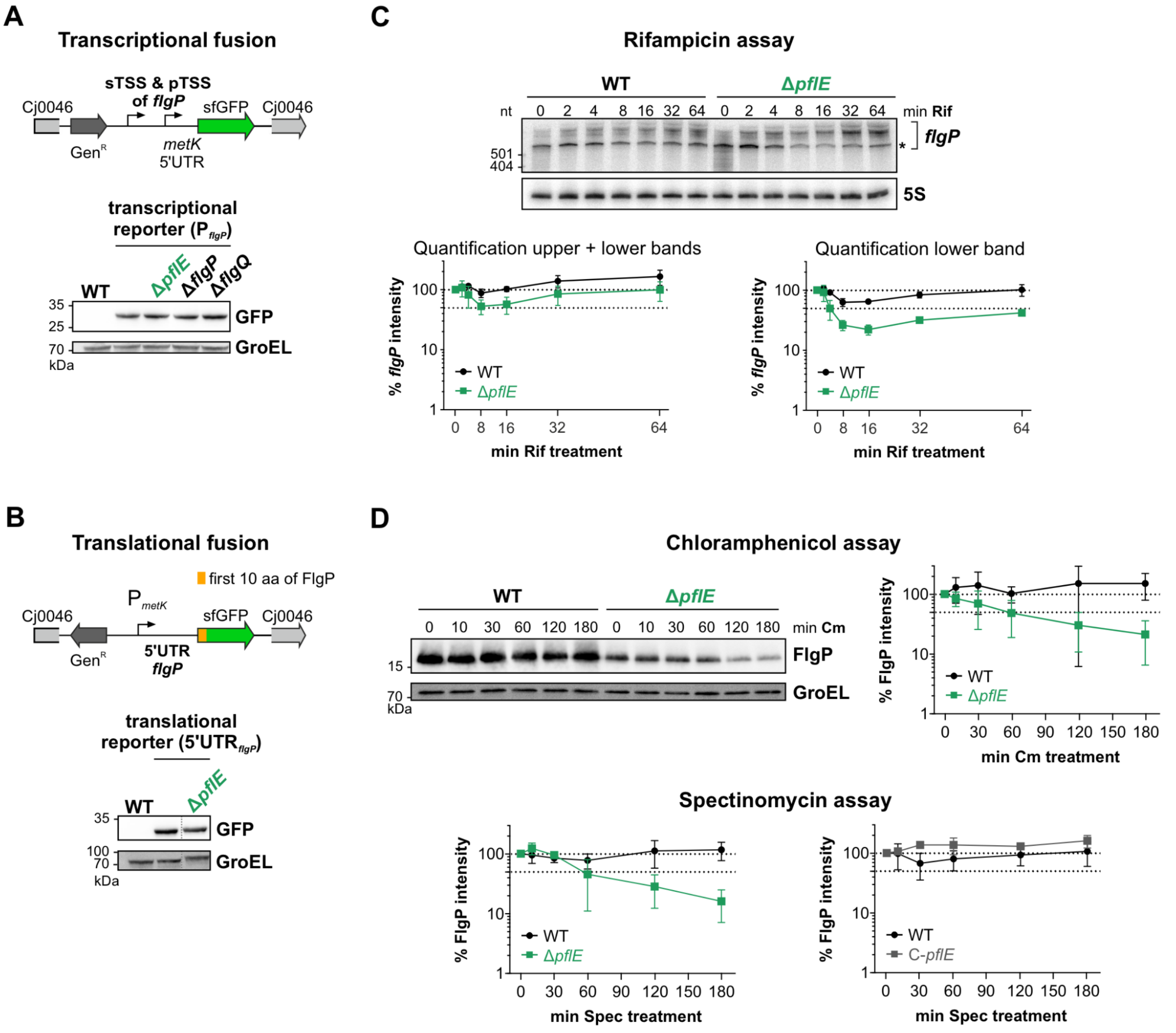


Figure S6. **PflE does not affect *flgP* transcription or translation but affects FlgP protein stability. (A)** Transcriptional reporter fusion of the *flgP* promoter to sfGFP. A potential secondary TSS (sTSS) as well as the primary TSS (pTSS) annotated for *flgP* (Dugar et al., 2013) were included in the 548 bp promoter fragment. The reporter was integrated in the *Cj0046* pseudogene locus and combined with the respective deletion mutants. Expression of the transcriptional reporter was examined by western blot analysis with an anti-GFP antibody. GroEL served as loading control. **(B)** Translational sfGFP reporter fusion of *flgP*. The reporter is transcribed from the constitutive $P_{metK}$ promoter, which was fused to the 27 nt 5'UTR of *flgP* (pTSS). The first 10 codons of *flgP* were fused to the second codon of sfGFP. As above, reporter expression was detected on a western blot with an anti-GFP antibody. GroEL served as loading control. Blots in panels A and B are representative of three biological replicates. **(C)** Rifampicin assay to assess mRNA transcript stability in WT and *pflE* deletion mutant. (*Left*) RNA was probed for *flgP* transcripts (CSO-5601) as well as 5S rRNA (CSO-0192) as a loading control. (*Right*) Quantification of rifampicin assay for two biological replicates, normalized to 5S rRNA signal. Either all bands above 500nt were quantified together (bracket & left graph) or the lower band only (asterisk & right graph). **(D)** Stability of FlgP protein levels in WT, Δ*pflE,* and C-*pflE.* Protein synthesis was halted with chloramphenicol (upper) or spectinomycin (lower), and samples were removed for western blot analysis with a FlgP antiserum at the indicated time points. The depicted % FlgP intensity represents the mean of n = 3 independent replicates with SD and is normalized to the loading control GroEL.

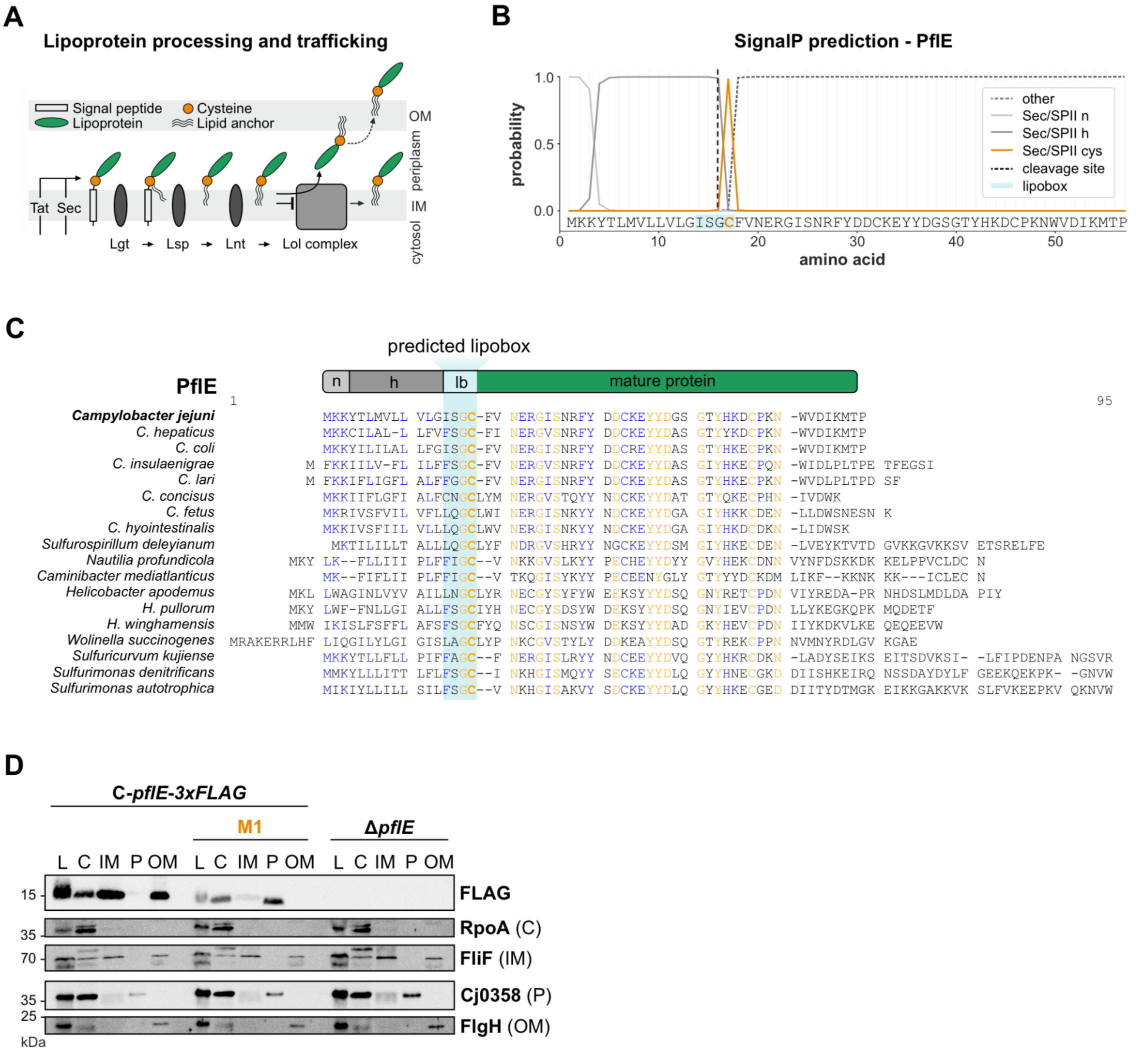


**Figure S7. PflE lipoprotein properties, conservation, and subcellular localization. (A)** Overview of lipoprotein processing and trafficking in bacteria. Adapted from: (Grabowicz, 2019) and (El Rayes et al., 2021). Tat: twin-arginine translocation machinery; Sec: general secretory system; Lgt: diacylglyceryl transferase; Lsp: lipoprotein signal peptidase; Lnt: apolipoprotein N-acyltransferase; IM: inner membrane; OM: outer membrane. **(B)** Signal peptide prediction of PflE with SignalP – 6.0 (Teufel et al., 2022). **(C)** Sequence alignment of PflE homologs from indicated Campylobacterota. Predicted lipoboxes are highlighted in light blue with the invariant cysteine marked in bold. Alignment was generated with Multalin (Corpet, 1988). n: N-terminal region. H: hydrophobic region. lb: lipobox. **(D)** Subcellular localization of PflE-3xFLAG in complementation strains complemented with PflE wild-type construct or the invariant cysteine mutant C17A (M1) as well as in Δ*pflE*. Strains were subjected to subcellular fractionation and analyzed by western blot. PflE-3xFLAG was detected by anti-FLAG. L: lysate; C: cytoplasm; IM: inner membrane; P: periplasm; OM: outer membrane. Controls for fractionation: (C) cytosolic protein RpoA (anti-RpoA; UTGP275; (Waller et al., 2024)), (IM) inner membrane protein FliF (anti-FliF; M202; (Boll and Hendrixson, 2013)), (P) periplasmic protein Cj0358 (anti-CJJ81176_0382; M17; (Bingham-Ramos and Hendrixson, 2008)), and (OM) outer membrane protein FlgH (anti-FlgH; UTGP161).

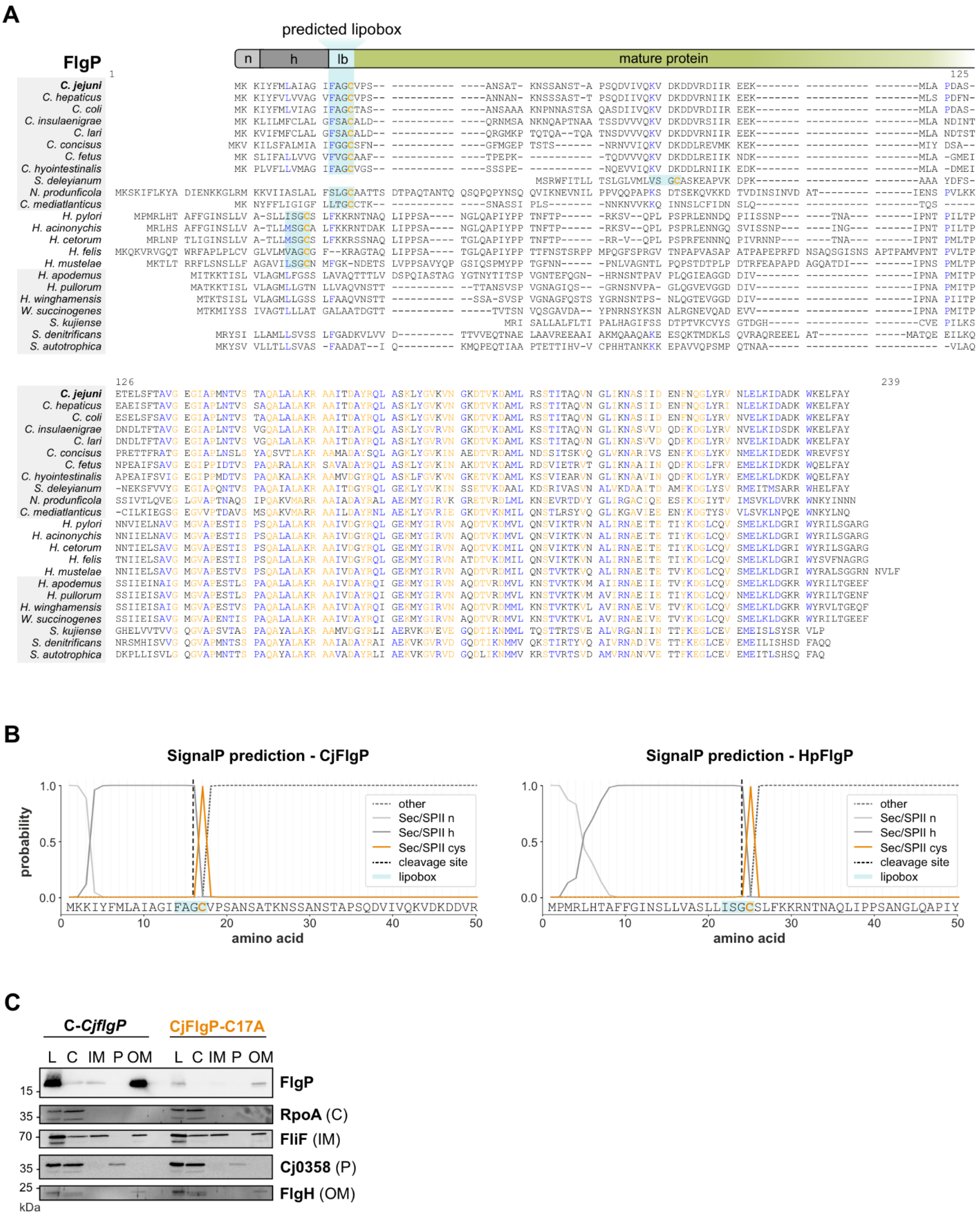

**Figure S8. FlgP conservation, lipoprotein properties, and subcellular localization. (A)** FlgP homologs were detected by blastP using default parameters or the orthologue function at KEGG. Lipoprotein signal peptides/cleavage sites were detected with SignalP v6 (probability >0.9) (Teufel et al., 2022) using default parameters. Putative lipoboxes, including the invariant Cys, are highlighted in light blue and in bold and orange, respectively. Grey: Species conserving *pflE*. n: N-terminal region. H: hydrophobic region. lb: lipobox. **(B)** Signal peptide prediction of FlgP with SignalP – 6.0 (Teufel et al., 2022) in *C. jejuni* (CjFlgP, left) and *H. pylori* (HpFlgP, right). **(C)** Subcellular localization of FlgP in a Δ*flgP* mutant complemented with wild-type FlgP (C-*CjflgP*) or the lipobox mutant of FlgP (CjFlgP-C17A) in the *rdxA* locus. Strains were subjected to subcellular fractionation and analyzed by western blot. FlgP was detected by FlgP antiserum. L: lysate; C: cytoplasm; IM: inner membrane; P: periplasm; OM: outer membrane. Controls for fractionation: (C) cytosolic protein RpoA (anti-RpoA; UTGP275; (Waller et al., 2024)), (IM) inner membrane protein FliF (anti-FliF; M202; (Boll and Hendrixson, 2013)), (P) periplasmic protein Cj0358 (anti-CJJ81176_0382; M17; (Bingham-Ramos and Hendrixson, 2008)), and (OM) outer membrane protein FlgH (anti-FlgH; UTGP161).

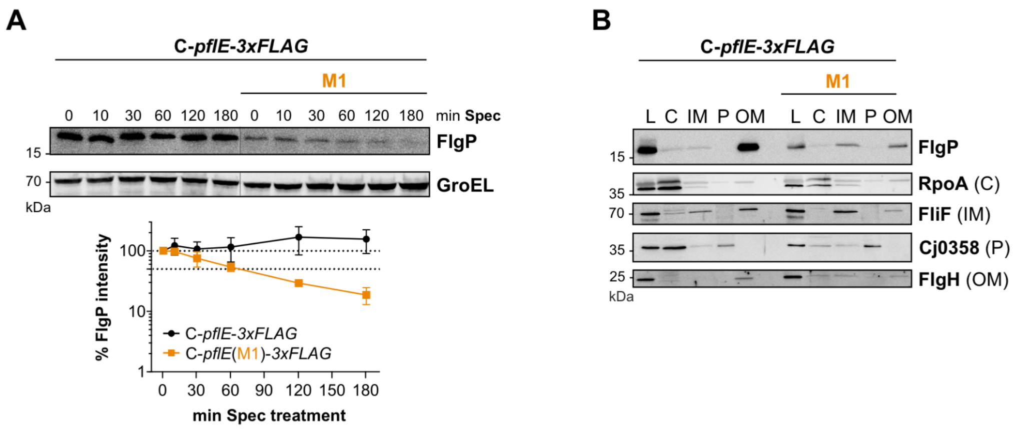


**Figure S9. FlgP protein stability and subcellular localization in PflE-M1. (A)** Protein stability assay for FlgP in *C. jejuni* NCTC11168 PflE complementation strains, either complemented with wild-type PflE sequence (C-*pflE-3xFLAG*) or the PflE-M1 lipobox mutant (M1) using the translation inhibitor spectinomycin (Spec). FlgP was detected with FlgP antiserum. GroEL: loading control. Quantification (lower) was performed with AIDA software, normalized to GroEL, and depicts the mean and standard deviation of independent replicates (n = 3). **(B)** Subcellular localization of FlgP in the same complementation strains as in (A). Strains were subjected to subcellular fractionation and analyzed by western blot. FlgP was detected by FlgP antiserum. L: lysate; C: cytoplasm; IM: inner membrane; P: periplasm; OM: outer membrane. Controls for fractionation: (C) cytosolic protein RpoA (anti-RpoA; UTGP275; (Waller et al., 2024)), (IM) inner membrane protein FliF (anti-FliF; M202; (Boll and Hendrixson, 2013)), (P) periplasmic protein Cj0358 (anti-CJJ81176_0382; M17; (Bingham-Ramos and Hendrixson, 2008)), and (OM) outer membrane protein FlgH (anti-FlgH; UTGP161).

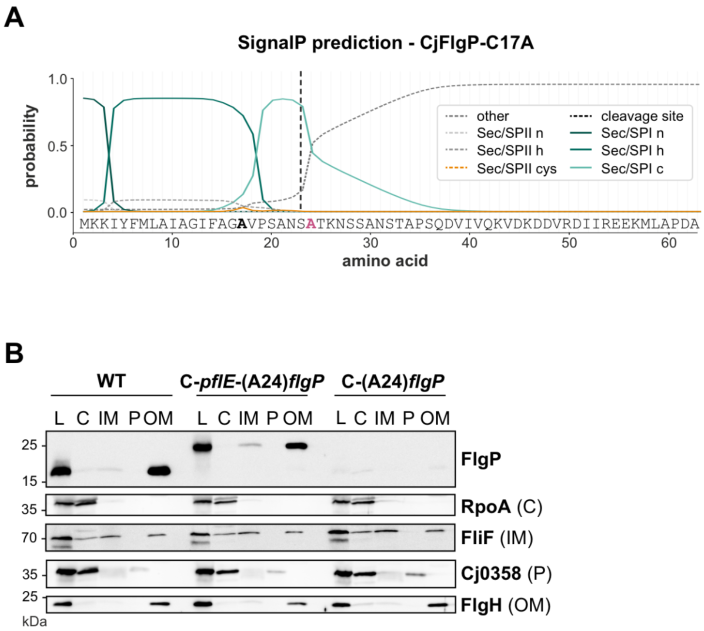


**Figure S10. Lipoprotein properties of CjFlgP-C17A and subcellular localization of PflE-FlgP fusion proteins. (A)** Signal peptide prediction of the FlgP lipobox mutant (CjFlgP-C17A) with SignalP – 6.0 (Teufel et al., 2022). C17A is highlighted in bold. Pink: predicted first amino acid of mature protein. n: N-terminus; h: helix; c: C-terminus; cys: invariant Cys of lipobox. **(B)** Subcellular fractionation of *C. jejuni* WT and a double deletion of *flgP* and *pflE* complemented with a *pflE-flgP* fusion construct (C-*pflE*-(A24)*flgP*) or a truncated version of FlgP (C-(A24)*flgP*). FlgP was detected with FlgP antiserum. L: lysate; C: cytoplasm; IM: inner membrane; P: periplasm; OM: outer membrane. Controls for fractionation: (C) cytosolic protein RpoA (anti-RpoA; UTGP275; (Waller et al., 2024)), (IM) inner membrane protein FliF (anti-FliF; M202; (Boll and Hendrixson, 2013)), (P) periplasmic protein Cj0358 (anti-CJJ81176_0382; M17; (Bingham-Ramos and Hendrixson, 2008)), and (OM) outer membrane protein FlgH (anti-FlgH; UTGP161).
